# Heterogeneous mutation rates and spectra in yeast hybrids

**DOI:** 10.1101/2021.06.04.447117

**Authors:** Anna Fijarczyk, Mathieu Hénault, Souhir Marsit, Guillaume Charron, Christian R. Landry

## Abstract

Mutation rates and spectra vary between species and among populations. Hybridization can contribute to this variation, but its role remains poorly understood. Estimating mutation rates requires controlled conditions where the effect of natural selection can be minimized. One way to achieve this is through mutation accumulation experiments coupled with genome sequencing. Here we investigate 400 mutation accumulation lines initiated from 11 genotypes spanning intra-lineage, inter-lineage and interspecific crosses of the yeasts *Saccharomyces paradoxus* and *S. cerevisiae* and propagated for 770 generations. We find significant differences in mutation rates and spectra among crosses, which are not related to the level of divergence of parental strains but are specific to some genotype combinations. Differences in number of generations and departures from neutrality play a minor role, whereas polyploidy and loss of heterozygosity impact mutation rates in some of the hybrid crosses in an opposite way.

## Introduction

Mutations generate genome variation, which in turn fuels evolution. They can be a source of adaptations (Venkataram et al. 2016), though more often they contribute to increased genetic load and play an important role in disease development (Deciphering Developmental Disorders Study 2017), in particular the evolution of cancer (ICGC/TCGA Pan-Cancer Analysis of Whole Genomes Consortium 2020). Mutations result from DNA replication errors and DNA damage, from the activity of transposable elements and other physical alterations of DNA molecules. The balance between mechanisms generating and diminishing mutations results in largely conserved mutation rates within large taxonomic units (Drake 1991; Lynch 2007), but to what extent rates of mutations and different mutation types (mutation spectra) differ among populations and between closely related species is only beginning to be understood.

Whole genome population data can provide estimates of mutation rates; however, whether novel mutations become observable variants in natural populations depends on environmental variation, selection, recombination, and demographic history. Largely unbiased mutation rates can be measured in microorganisms using fluctuation assays (Luria and Delbrück 1943) or mutation accumulation (MA) experiments followed by reporter assays or genome sequencing. In MA experiments, the strains are first propagated through sequential bottlenecks for many generations and then sequenced to detect mutations. Both approaches have been used in different contexts with model organisms. In yeast, mutation rates and spectra have been shown to differ within and between species (Nguyen et al. 2020; Jiang et al. 2021), between environmental factors such as mild (Liu and Zhang 2019) and severe stressors (Shor et al. 2013) and vary depending on ploidy (Sharp et al. 2018) or sequence context (Ma et al. 2012). A wide array of genes, notably involved in DNA repair, have been shown to influence the mutation landscape (Demogines et al. 2008; Lang et al. 2013; Serero et al. 2014; Stirling et al. 2014; Gou et al. 2019; Loeillet et al. 2020).

Hybridization could play a role in shaping mutation rates as it could compound the effects of different mutator alleles (Demogines et al. 2008), introduce mutations through mutagenic crossing-over in high diversity regions (Yang et al. 2015) or disrupt mechanisms repressing transposable elements (Petrov et al. 1995; Serrato-Capuchina and Matute 2018). Alternatively, parental effects can dominate in shaping mutation rates in hybrids (Bashir et al. 2014). MA experiments by Tattini et al (Tattini et al. 2019) showed that an interspecific hybrid between one *S. cerevisiae* and *S. paradoxus* strain has mutation rates similar to *S. cerevisiae* diploid strain. However, to better understand the role of hybridization and genotype effects on mutation rate, different genetic backgrounds need to be considered.

Here we analyzed whole genome sequences from our previous MA experiments (Charron et al. 2019; Hénault et al. 2020; Marsit et al. 2021) to estimate and compare mutation rates and spectra in nine intraspecific crosses of *Saccharomyces paradoxus* and two interspecific crosses between *S. paradoxus* and *S. cerevisiae* yeast. We find that parental sequence divergence does not explain differences in mutation rate, whereas polyploidy and loss of heterozygosity (LOH) correlate negatively or positively with mutation rate. Our results suggest that combinations of individual genotypes best determine mutation rates and spectra among crosses.

## Results and Discussion

### Mutation rate and spectrum

We analyzed 400 lines from 11 crosses, including 5 intra-lineage crosses of *S. paradoxus SpC* (CC) and *SpB* (BB), 4 inter-lineage crosses between *S. paradoxus SpB* and *SpC* (BC) and between *SpB* and *SpA* (BA) and 2 interspecific crosses between *S. paradoxus SpB* and *S. cerevisiae* (BSc) (fig. 1A,B, supplementary tables S1-4). After ∼770 generations of mutation accumulation (MA), lines accumulated a total of 1040 single and multiple nucleotide substitutions and 34 indels (supplementary tables S5-6). Mean per-cross mutation rates range from 7.88×10^−11^ in the inter-lineage cross BA2 (4% parental nucleotide divergence) to 2.64×10^−10^ in the intra-lineage cross BB2 (0.3% parental nucleotide divergence) and they are generally consistent with mutation rates found previously in both laboratory and natural strains of *S. cerevisiae, S. paradoxus* and their hybrids (fig. 1C, supplementary table S7). Mutation rates vary significantly among crosses (negative binomial regression, *P*-value = 1.11×10^−6^, supplementary table 8). Mutation rates also vary between crosses which share one of their parental strains (Kruskal-Wallis test, *P*-values 0.025 and 7.24×10^−5^ for B1 and B2 groups respectively), and the trends are not monotonic with sequence divergence of parental strains (fig. 1C). Differences in mutation rates between pairs of crosses of the same lineage are generally lower, but of the same order of magnitude than those between pairs of crosses from different lineages, with the highest differences for pairs including BB crosses (supplementary fig. 1).

**Fig. 1.**
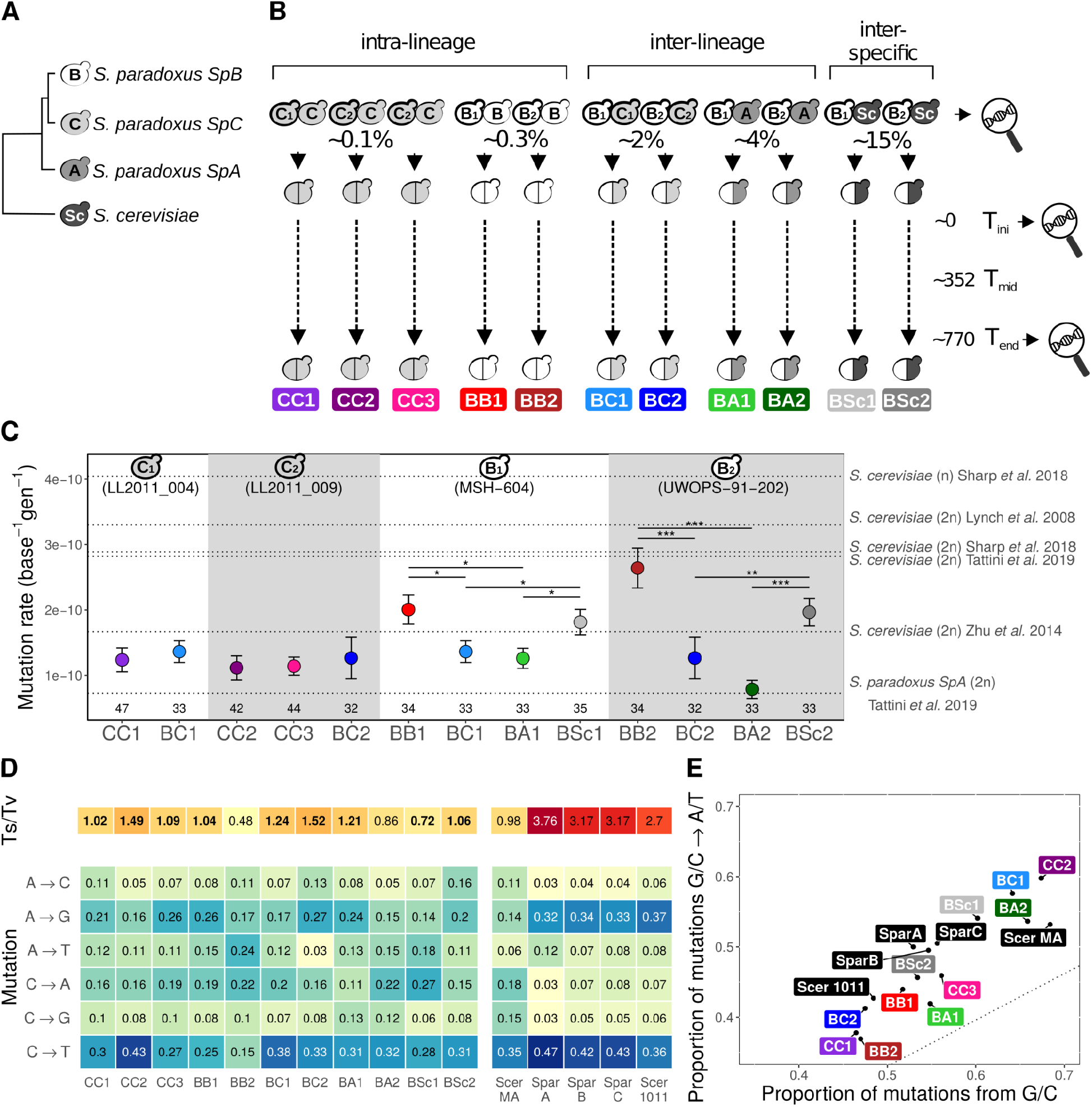
Mutation rates differ among crosses. **A**. Schematic phylogenetic relationships among the lineages. **B**. Analyzed crosses. Two *S. paradoxus SpB* and two *S. paradoxus SpC* parental strains that were used in multiple crosses are marked with numbers and bold contour (shown in panel C and detailed in supplementary table S1). Parental strains without numbers correspond to distinct strains. Percentages under the crosses indicate % of sequence divergence between parental genomes. Whole genome sequencing was done in parental strains, at T_ini_ and T_end_ and growth was measured at three timepoints (T_ini_, T_mid_, T_end_). **C**. Mean mutation rates per haploid position per generation with standard errors estimated from replicate lines over 770 generations of MA experiment shown in four groups sharing the same parental strain. Asterisks show FDR-corrected differences at \**P*-value < 0.1,\*\**P*-value < 0.01, \*\*\**P*-value < 0.001, Wilcoxon Rank Sum Test. Number of lines is depicted at the bottom. **D**. Transition to transversion ratio (Ts/Tv, upper red heatmap) and frequencies of 6 nucleotide changes including their complementary changes (lower blue heatmap). Scer MA stands for mutation spectrum of *S. cerevisiae* from the mutation accumulation experiment from (Zhu et al. 2014). The three Spar columns correspond to a population dataset of 3 lineages of *S. paradoxus*: *SpA, SpB* and *SpC*. Scer 1011 corresponds to the population dataset from (Peter et al. 2018). Bold Ts/Tv estimates indicate crosses with Ts/Tv significantly different from 0.5 (chi-square contingency test, FDR-corrected *P*-value < 0.01). **E**. All crosses show higher frequency of G/C to A/T than expected from the proportion of mutated G/C positions. The dotted line represents the expected proportion of mutations G/C to A/T if mutations were random.

There are small but nonsignificant differences (chi-square contingency test, FDR-corrected *P*-value > 0.98) in frequency spectra of 6 mutation types (fig. 1D, after excluding 2 lines, see Methods). Mutations of type C>T (G>A) are most frequent in all crosses except BB2, which also has the lowest transition to transversion (Ts/Tv) ratio close to 0.5 (fig. 1D). Unlike *S. cerevisiae*, most *S. paradoxus* crosses show in general a high frequency of A>G (T>C) mutations, which results in a smaller fraction of G/C positions being hit by a mutation (fig. 1E). Similar to *S. cerevisiae*, we observe a bias of G/C to T/A mutations in hybrid crosses (fig. 1E). Again, we find no differences between intra- and inter-lineage crosses, and BB2 stands out in terms of both mutation rate and spectrum.

We investigated several biological factors which could explain differences in mutation rate and spectra among crosses and lines including differences in number of generations, deviations from neutral evolution, whole genome or chromosome ploidy changes, the presence of loss of heterozygosity (LOH) events and disruption of DNA repair genes.

### Number of generations

Mutation rate estimates were calculated with an average estimation of the number of mitotic divisions within a colony during the MA experiments (Charron et al. 2019). However, these estimates can be biased by variation in actual generation time. We sought to systematically correct for this by measuring the growth of all individual lines at three timepoints of the MA experiments by extracting the maximal yield from growth curves in liquid medium (fig. 2A), and using these values as a proxy of generation time (supplementary figure S2). Overall, growth yields showed no significant trend with timepoints (mixed linear model with cross-specific intercepts and slopes, *P*-value = 0.93), which is consistent with the absence of selection for growth efficiency. There are significant differences between crosses and timepoints (fig. 2A, two-way ANOVA, *P*-values = 5.7×10^−23^ and 8×10^−5^ respectively), but no significant interaction between crosses and timepoints (*P*-value = 0.56). We find that growth yield does not explain differences in mutation rates between crosses. Neither mean yield (across three timepoints) nor mean growth yield by cross interaction have a significant effect on mutation rate (fig. 2B, supplementary figs S3-4, supplementary table S9, negative binomial regression, *P*-values > 0.718 for both growth and interaction).

**Fig. 2.**
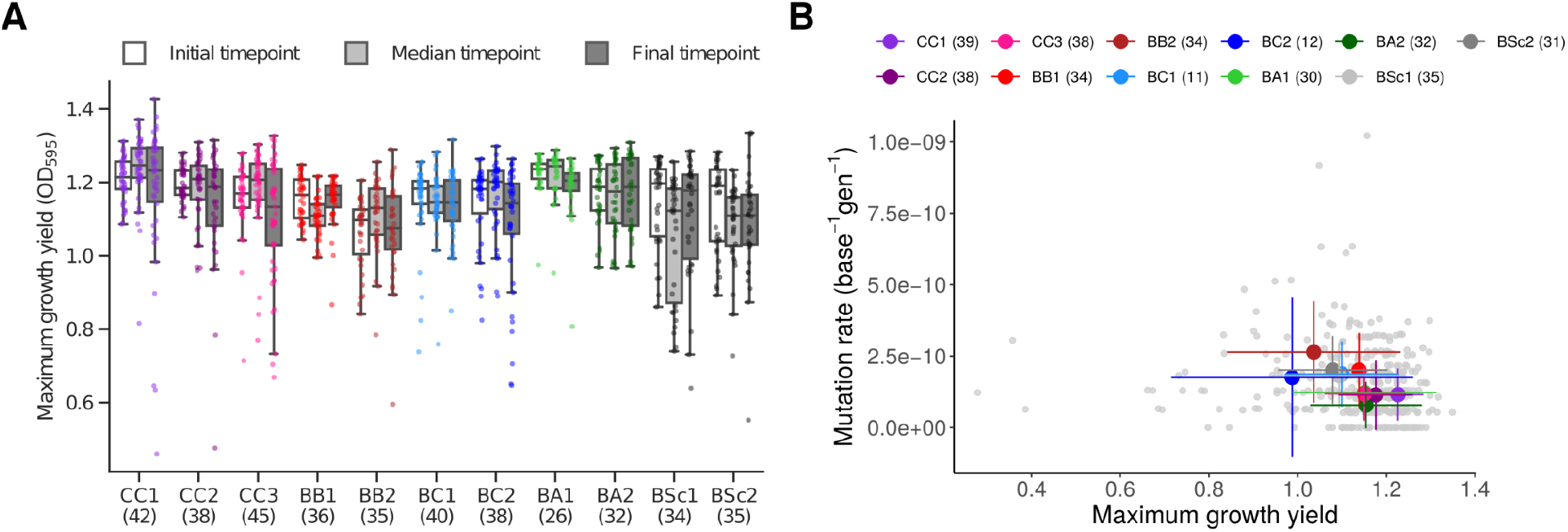
Number of generations does not explain mutation rate variation. **A**. Growth yields differ significantly between crosses. The maximum growth yield of individual lines at the initial, median, and final timepoints of the MA experiment are shown with sample sizes in parentheses. **B**. Mean growth rate (across three timepoints per cross) vs. mean mutation rates per cross with errors depicting standard deviation across lines (diploids only) with sample sizes in parentheses.

### Functional properties of *de novo* mutations

The demographic regimen of MA experiments is designed to minimize the efficiency of natural selection on the fixation of spontaneous mutations, which should result in fixed mutations that are randomly distributed in genomes, assuming a uniform mutation rate. We tested this hypothesis by classifying the *de novo* mutations according to three functional properties expected to impact their average fitness effect: whether they occur in protein-coding sequences, and if so, whether they lead to non-synonymous substitutions or fall into essential genes. Assuming that spontaneous mutations are deleterious on average, efficient purifying selection would leave fewer mutations than expected by chance in coding regions and essential genes, and non-synonymous mutations should be rare. We examined these predictions while accounting for the genomic composition and substitution spectrum of each cross (supplementary figs. S5-8; supplementary table S10). Only BA1 exhibits a pattern consistent with purifying selection, with more mutations in non-essential genes than expected. In the BA1 lines, only three mutations were identified in essential genes. Despite all three being non-synonymous, they are not associated with significant growth yield decrease (supplementary fig. S9). Although growth differences may fall below our method’s detection limit, this suggests that the scarcity of mutations in essential genes is not linked to increased fitness. On the other hand, four crosses exhibit opposite patterns for at least one property, with less non-coding mutations (BB1, BSc1), less mutations in non-essential genes (BB2) and more non-synonymous mutations (BB2, BA1). Nevertheless, no pattern was consistent with increased growth yield (supplementary fig. S9). Overall, these results indicate that accumulated mutations are largely consistent with neutral evolution.

### Polyploidy and loss of heterozygosity

It was shown that *S. cerevisiae* diploids have lower mutation rates than haploids (Sharp et al. 2018). One type of change that can occur in diploids is LOH. LOH can eliminate arising *de novo* mutations, but on the other hand, it is an error-prone process (Hicks et al. 2010; Deem et al. 2011), which can lead to higher mutation rates. We obtained LOH tracts from (Marsit et al. 2021) for diploid and triploid lines from crosses BB, BC, BA and BSc. To include CC crosses and LOH events smaller than the minimum LOH tract length (1 kb) we additionally calculated LOH rate using the proportion of heterozygous positions in which genotype changed between T_ini_ and T_end_ (supplementary figs. S10-11). We find a significant negative effect of polyploidy and a significant positive effect of LOH on mutation rate (supplementary table S11). The effect of interaction between polyploidy and LOH is negative but nonsignificant (negative binomial regression, *P*-value = 0.08). In BC crosses, which carry most triploid lines, BC1 triploids have significantly lower mutation rates than BC1 diploids, whereas in BC2 there is no difference (fig. 3A). We confirmed that differences are not caused by uncalled mutations in low covered regions (supplementary fig. S12). LOH rates are slightly higher for triploids than diploids, though the difference is nonsignificant (fig. 3B). Large scale LOH events (> 300 bp) are less frequent in BC triploids than diploids (Marsit et al. 2021, supplementary fig. 13), suggesting that total observed LOH rates in triploids are driven mostly by very short LOH events, which do not disappear after excluding positions below read depth of 70x (supplementary fig. 12). This implies that LOH events do not have a detectable impact on lowering mutation rates in polyploids at this stage of experiment. Contrary to triploidy, gains or losses of chromosomes, and whole genome duplications occuring during the experiment do not have a measurable effect on mutation rate (supplementary fig. S14).

**Fig. 3.**
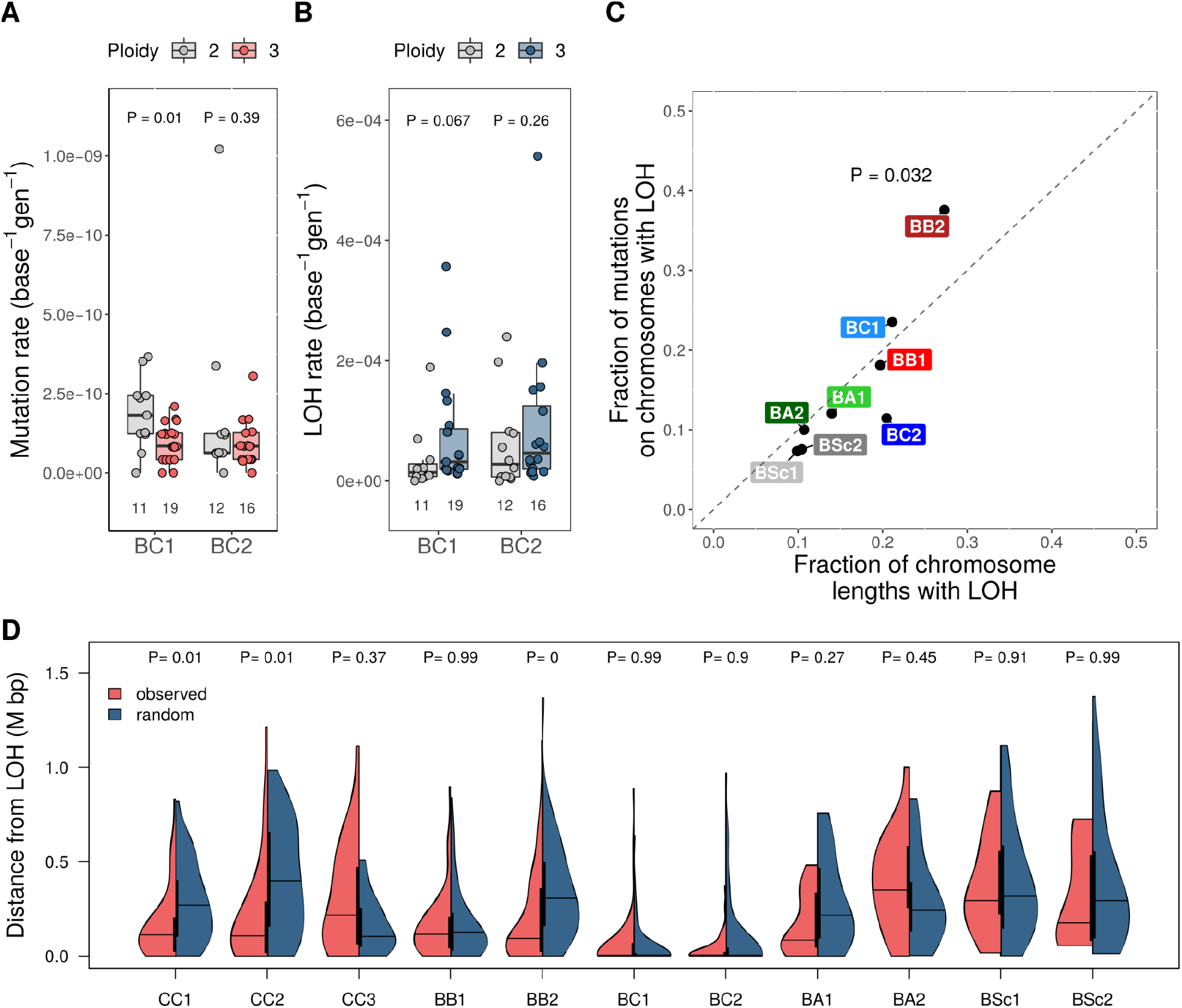
Polyploidy and LOH impact mutation rates. **A**. Differences in mutation rates in BC crosses between diploid and triploid lines tested with Wilcoxon rank sum test. **B**. Differences in LOH rates in BC crosses between diploid and triploid lines tested with Wilcoxon rank sum test. **C**. Fraction of mutations on chromosomes with LOH compared to expected (diagonal), estimated as the fraction of the genome comprising the sum of chromosome lengths with LOH. BB2 has significantly more mutations on LOH-carrying chromosomes than expected from the length of LOH-carrying chromosomes. **D**. Distribution of distances of *de novo* mutations to the closest LOH variants compared with the same number of mutations randomly drawn from the genome. Differences compared with Wilcoxon rank sum test.

We then investigated whether LOH can induce mutations in diploid lines. Chromosomes carrying LOH are not enriched for *de novo* mutations, except in BB2 (binomial test, fig. 3C). We find no difference in the number of *de novo* mutations around the breaks of LOH segments (+/-2 kbp) compared to the remainder of chromosomes carrying LOH (binomial test, 2 out of 109 mutations, expected frequency = 0.012, *P*-value = 0.395), which is not unexpected due to ambiguous LOH break positions for low heterozygosity crosses. However, *de novo* mutations in BB2, CC1 and CC2, are significantly closer to LOH positions than random mutations, suggesting the causal effect of LOH on mutation rate (fig. 3D). Increased mutation rates near LOH breakpoints were observed before (for example, in pathogenic *Candida* yeast (Ene et al. 2018)), which can be linked to mutagenic effects of homologous recombination. Even though the duration of our experiments is relatively short to observe strong LOH effects, our results suggest that some genotype combinations may experience stronger mutagenic effects of homologous recombination.

### DNA repair genes

Observed differences in mutation rates and spectra are mainly driven by BB2, which has the highest mutation rate and the lowest Ts/Tv ratio. Malfunctioning DNA repair mechanisms can influence both mutation rates and spectra. One of the patterns observed in yeast with a disrupted DNA mismatch repair pathway is the tendency to accumulate spontaneous mutations in regions with repeated sequence motifs (Lang et al. 2013). BB crosses have indeed the largest (although nonsignificant) departure from the expected sequence complexity around *de novo* mutations towards lower values (supplementary fig. S15). Disruption of DNA repair genes with an impact on all lines from the cross could occur before the formation of the cross or before sequencing at T_ini_. We found no common non-synonymous or non-sense variants at the beginning of MA in DNA repair genes in BB2. Also none of the DNA repair genes lost a copy (supplementary fig. S16), and lines with lost, gained, or unbalanced number of copies of DNA repair genes do not have an increased mutation rate (supplementary fig. S15). On the contrary, outlier lines with the highest mutation rates in five crosses have a non-synonymous mutation in at least one DNA repair gene (supplementary fig. S17). Presence of *de novo* non-synonymous or mutations causing stop codon gain/loss in DNA repair genes is a strong predictor of mutation count (negative binomial regression, *P*-value = 8.79e-8, incidence rate ratio = 2.6, CI: 1.8-3.7). Acquired mutations, therefore, explain part of the variation in the mutational landscape. Skewed mutation rate and spectrum in BB2 is not driven by a single line but elevated A>T and C>A mutations across multiple lines, most often surrounded by A and T nucleotides (supplementary fig. 18, supplementary table S5). This indicates that parent-specific factors or between parent interactions must account for this change. The list of 22 DNA repair genes with non-synonymous private variants in the parental strain specific to BB2 (LL2012_021, supplementary table S12), includes a gene encoding DNA mismatch protein MSH2, which binds to DNA mismatches upon heterodimerization with MSH6 (Marsischky et al. 1996). Mutation Lys957Glu is located in a conserved position (ConSurf conservation score = -0.697, CI: -1.022 to -0.501 (Landau et al. 2005)) in the region of dimerization with MSH6, pointing to a potential trigger of biased mutational landscape in BB2 cross.

In this study we demonstrate significant differences in mutation rates between 11 types of hybrid crosses of yeast. Including genetically diverse backgrounds is therefore essential for uncovering species diversity in mutation rates. Low mutation rates in *S. paradoxus SpA* reported previously (Tattini et al. 2019) are closest to BA2 (4% parental divergence), which has the lowest mutation rate among all crosses. Out of several potential factors leading to such variation, differences in growth rate and natural selection play a negligible role in our experiment. Polyploidy and LOH have observable and opposite impacts on mutation rates of some crosses. LOH is generally more frequent in less heterozygous hybrids (Tattini et al. 2019; Marsit et al. 2021), therefore its effects should be more prominent in crosses involving closely related lineages (BB and CC). On the contrary, the level of sequence divergence between parental strains does not have a systematic impact on mutation rate. This is consistent with the results of (Hénault et al. 2020) who found no relationship between the accumulation of transposable elements and the level of divergence between parental genomes in the same crosses. Even though we are not able to measure the impact of each parental genotype on mutation rate in this experimental setup, our results suggest that hybrid genotypes show variation of mutation rates and spectra either through inheritance of parent-specific rates or interaction between parental genotypes. In consequence we may expect the natural populations of *S. paradoxus*, which has experienced many hybridization events in the recent past (Leducq et al. 2016; Eberlein et al. 2019), to exhibit such variation.

## Materials & Methods

### Hybrid crosses, mutation accumulation experiments and genome sequencing

The detailed methods for the generation of hybrid crosses, the mutation accumulation experiments and the whole-genome sequencing are described in (Charron et al. 2019; Hénault et al. 2020; Marsit et al. 2021). Briefly, hybrids were generated by crossing haploid strains of *S. paradoxus* or *S. cerevisiae* of genotype MATa or MATα, *ade2Δ*::hphNT1, and *hoΔ*::kanMX or *hoΔ*::natMX, and by selecting for dual resistance to geneticin and nourseothricin. From 48 to 96 lines (each derived from independent mating events) were initiated for each of a set of 11 different crosses, spanning a range of genetic divergence levels: within *S. paradoxus SpC* (CC1, CC2, CC3); within *S. paradoxus SpB* (BB1, BB2); between *S. paradoxus SpB* and *SpC* (BC1, BC2); between *S. paradoxus SpB* and *SpA* (BA1, BA2); and between *S. paradoxus SpB* and *S. cerevisiae* (BSc1, Bsc2). All the lines were evolved at room temperature on YPD solid medium (1% yeast extract, 2% tryptone, 2% dextrose, 2% agar) for 35 passages. Each passage was performed by streaking one randomly selected single colony (the closest to a predefined position on the plate) for single colonies every three days, thus creating periodic single-cell bottlenecks in each line. Glycerol stocks of all the lines were sampled at intervals of three passages. From 23 to 40 lines were selected for whole-genome sequencing of the stocks at the initial and final timepoints. Shotgun genomic DNA libraries were prepared with the Illumina Nextera kit and a modified protocol (Baym et al. 2015). Libraries were sequenced in paired-end 150 bp mode on HiSeqX or NovaSeq 6000 systems. Short read sequence data is available from NCBI under accession PRJNA515073.

### Growth rate measurements

Maximal growth rates in liquid cultures were measured to approximate the number of generations for which each MA line was evolved in the evolution experiments. For each sequenced line, the stocks from the initial, median and final timepoints were selected for growth measurement. Growth measurements were performed following a randomized design. Two technical replicates of each line at each timepoint were included. For each combination of (cross, timepoint), all lines were randomly assigned to one of seven measurement experiments. Within each experiment, lines and replicates were randomly assigned to a position in a 16 by 24 array. Glycerol stocks were thawed and spotted on YPD solid medium (1% yeast extract, 2% tryptone, 2% dextrose, 2% agar). Plates were incubated for 72 hrs at room temperature. Precultures were inoculated in 0.5 ml of YPD liquid medium and incubated at room temperature for 24 hrs. 50 μl of preculture was diluted in 1 ml of fresh YPD liquid medium. Two 80 μl aliquots of this dilution were distributed into sterile 384-well polystyrene microplates (BRANDplates pureGrade S 384, BRAND GMBH + CO KG, Wertheim, Germany) using an Eppendorf epMotion 5075 liquid handling robotic platform (Eppendorf, Hamburg, Germany). Incubation at a controlled temperature of 25°C and optical density (OD_595_) measurements at 15 min intervals for 48 hrs were performed in a Tecan Spark microplate reader (Tecan Group Ltd, Männedorf, Switzerland) equipped with a cooling module. Replicates with no growth or abnormally high initial OD_595_ values were discarded. Values were normalized by subtracting the timewise average of empty wells. Maximum growth yield of each line at each timepoint of the experiment was measured as the maximum OD_595_ (averaged over the two technical replicates) reached within the 48 hours of growth measurement. Maximum growth rate was inferred by fitting a Gaussian process to the curves, using the empirical variance of replicates to estimate technical error (Swain et al. 2016).

To assess the validity of growth in liquid medium as a proxy of colony growth on solid medium, we measured cell counts for colonies propagated using the same method as for MA lines propagation. Glycerol stocks for 24 randomly selected MA lines (all crosses and timepoints confounded) were spotted on YPD solid medium and incubated at room temperature for three days. Spots were streaked for single colonies on YPD solid medium on thirds of 10 mm Petri dishes and incubated for three days at room temperature. Using the same procedure, four single colonies per line were streaked. Single colonies from the second streakings were extracted by cutting agar cubes with a sterile scalpel blade. For each line, agar cubes were added to 750 μl of sterile water in a 96-well deepwell plate, and resuspended by adding 750 μl of sterile water and mixing with gentle up-and-downs. 20 μl of resuspension were diluted in 180 μl of sterile water. Cell density was measured with a Guava easyCyte HT flow cytometer (Millipore Sigma, Burlington, USA). The OD_595_ of undiluted colony resuspensions was measured with Tecan Infinite F200 Pro microplate reader (Tecan Group Ltd, Männedorf, Switzerland).

### *De novo* mutation rates

All reads were trimmed for Illumina Nextera adapters using Trimmomatic v0.33 (Bolger et al. 2014) with options ILLUMINACLIP:nextera.fa:6:20:10 MINLEN:40. Reads from lines derived from *S. paradoxus* crosses were mapped to *S. paradoxus* assembly from the *SpB* lineage (MSH-604, crosses BB1, BB2, BC1, BC2, BA1 and BA2) or *SpC* lineage (LL2011_012, 3 *SpC* x *SpC* crosses CC1, CC2, CC3). Read mapping was done with bwa mem v0.7.17 (Li 2013). To minimize the effect of divergence between the parents of *S. cerevisiae* and *S. paradoxus* on mapping accuracy, reads coming from crosses BSc1 and BSc2 were mapped to concatenated assemblies of *S. paradoxus SpB* and *S. cerevisiae* (YPS128, (Yue et al. 2017)). Picard tools v2.18 (http://broadinstitute.github.io/picard/) were used to correct ReadGroup IDs and mark duplicate reads. Nanopore assemblies of *S. paradoxus* are available from NCBI under accession PRJNA514804.

Overall sample ploidy was taken from (Marsit et al. 2021). Chromosome copy number was determined using Control-FREEC v11.5 (Boeva et al. 2012), by estimating copy number in 250 bp non-overlapping windows, and choosing one supported by the majority of windows. Variants were called for each cross separately using Freebayes v1.3.1 (Garrison and Marth 2012) with a ploidy preset for each sample and chromosome. Options -q 20 (minimum base quality) --use-best-n-alleles 4 (evaluate only 4 best SNP alleles) --limit-coverage 20000 (maximum read depth of 20000) -F 0.02 (at least 0.02 reads supporting alternate allele) were used. Variants overlapping repeats, telomeres and centromeres were excluded. Variants were filtered based on given criteria: minimum variant quality of 30 (QUAL>30) and quality by the count of alternate alleles higher than 10 (QUAL/AO>10), each alternate allele supported by at least 3 reads on forward and 3 on reverse strand (SAF>3 & SAR>3), at least one read placed to the right and one read placed to the left of the alternate variant (RPL>1 & RPR>1) and mean mapping quality for alternate alleles vs. reference allele between 0.9 and 1.05 (0.9<MQM/MQMR<1.05). On the sample level, variant alleles had to be supported by at least 4 reads and all genotypes (including parents) had to be supported by at least 10 reads. Finally, *de novo* mutations had to be absent in both parents and absent at time T_ini_, but present in T_end_ in at most one sample from each cross.

14 samples were identified as contaminants with Principal Component Analysis and were subsequently discarded. In spite of stringent variant calling, we observed reads from different crosses (parents) mapping with high quality in very low frequencies. To exclude the possibility of calling variants from these reads, we identified all parental variants separately from crosses mapped to *SpB* (BB1, BB2, BC1, BC2, BA1, BA2, BSc1 and BSc2) or *SpC* (CC1, CC2, CC3) and did not consider these positions for *de novo* variant detection.

Crosses BSc1 and BSc2 were processed with the same criteria, with an exception that *de novo* mutations on either of the genomes were considered conditional on the absence of the mutation in T_ini_ and one parental strain only. Also, small copy number variants were excluded. To check if some of the *de novo* mutations were shared between the two genomes e.g. due to loss of heterozygosity, two reference genomes were aligned with each other using minimap2 v2.17-r941 (Li 2018) and paftools.js liftover options was used to identify corresponding positions of mutations. None of the aligned mutations were overlapping.

*De novo* mutation rate per line was estimated as the number of *de novo* mutations acquired between T_ini_ and T_end_, divided by the number of generations times the haploid genome size. The haploid genome size was summed across chromosomes where haploid chromosome length equaled length of an assembled chromosome excluding repeats, telomeres, centromeres, positions with coverage less than 10x for a line and positions with parental variants divided by the ploidy of the chromosome. For mutation rate calculation, ploidy per line was estimated as an average ploidy of samples from T_ini_ and T_end_. To adjust for the differences in generation number in each line, the average generation number (770) was multiplied by the ratio: growth yield/median growth yield, where growth yield is mean growth yield estimated over 3 timepoints, and median corresponds to median growth yield over all lines. This correction was restricted to diploid lines, as we found a significant effect of ploidy on the optical density of cells (supplementary fig. 2D).

Statistical tests were performed with R v3.6.3 or Python v3.6.8. All significance values are given after correction for multiple tests using Benjamini-Hochberg (BH) method (except in supplementary figure 17, where Bonferroni correction was applied). To test different factors on mutation rates, generalized linear models were fitted to mutation counts per line. Mutation counts per 2 × 10 Mbp genome were derived from per line mutation rates. Number of lines with 0 mutations were higher than expected by the model, giving an overdispersion parameter of 2.07. The high number of lines with 0 mutations could result from the conversion of mutation rates to mutation counts and stringent filtering criteria. To take overdispersion into account we used negative binomial regression with glm.nb() function from the MASS package in R to fit cross, growth rate, polyploidy and LOH rate. *P*-values for each factor or interaction were obtained with the chi-square test using anova() function in R, by comparing it with the simpler model lacking that factor.

### Loss of heterozygosity (LOH)

To estimate the impact of LOH on mutation rate we calculated LOH rate by counting how many heterozygous positions between parents changed genotype between T_ini_ to T_end_. In diploids it corresponds to the proportion of heterozygotes at T_ini_, which turned to homozygotes by T_end_. In triploids it includes cases of complete loss of heterozygosity (genotypes 0 and 1 for the parents, 0/0/1 (or 0/1/1) at T_ini_ and 0/0/0 (or 1/1/1) at T_end_) as well as change of genotype without the loss of heterozygosity (genotypes 0 and 1 for the parents, 0/0/1 (or 0/1/1) at T_ini_ and 0/1/1 (or 0/0/1) at T_end_). In aneuploid chromosomes (both in diploid and triploid lines), changes in genotypes were recognized as LOH only for positions having the same number of copies in T_ini_ and T_end_. Because each position is treated separately without the context of the surrounding genotypes, it is possible to include double LOH events in LOH rate calculation in triploids. This approach includes any LOH event, even those concerning a single position, and therefore can be applied to crosses with low heterozygosity, such as CC. To test whether this method can overestimate LOH rate in low covered regions, rates were estimated for subsets of data with the minimum coverage spanning from 10x to 150x. Triploid lines showed elevated rates for regions including low read depth, we therefore considered for final rate calculation only positions with the minimum read depth of 70x in T_ini_ and T_end_ for crosses CC, BB, BC and BA, and minimum 40x for crosses BSc. At the same time, we obtained LOH tracts from (Marsit et al. 2021), for crosses BB, BC, BA and BSc. As the goal of that study was to identify and quantify large scale LOH events (> 300 bp) the method for LOH detection was different. In short, heterozygous loci with allele frequencies deviating from the average allele frequency over the whole chromosome (±0.15) were kept as potential loci belonging to an LOH tract. Stretches of consecutive marker positions were grouped in LOH regions. Blocks of LOH were identified by looking for a minimum of three successive SNPs with the same allele frequency (±0.1) in a window of 300 bp for hybrids and a window of 1 kb for BB crosses. Because the minimum LOH size was of 1 kb in BB crosses due to the very low heterozygosity in these crosses, only LOH segments larger than 1 kb were considered for comparison between hybrids and BB crosses in this method (Marsit et al. 2021). LOH rates per bp per generation calculated with the two methods show generally consistent results with expected lower rates estimated from LOH tracts in low heterozygosity crosses, due to sparse heterozygous positions and LOH tract size limit (1kb) (supplementary fig. 11). To calculate the expected LOH rates from the rates based on LOH tracts, if no windows were missed due to low heterozygosity, observed rates were divided by the cross specific probability of LOH detection. Probability was calculated as the proportion of random 1000 bp windows outside of repeated regions, carrying at least 3 differences between parental genomes in BB crosses and at least 10 differences in BC, BA and BSc crosses.

### Mutation spectra

Inspection of mutation spectra across all experimental lines showed that 2 outlier lines with a substantial number of mutations have highly skewed mutation spectra. CC1_48 has 14 A>T and 7 A>G nucleotide substitutions out of a total of 22 and BC2_75 has 11 A>G mutations out of a total of 17 mutations. The two samples were excluded from the mutation spectra analyses.

To obtain mutation spectra for natural populations of *S. paradoxus* we took a population dataset of biallelic single nucleotide variants from (Eberlein et al. 2019). Genome of *S. cerevisiae* (YPS128) was aligned to determine the ancestral states. Hybrid lineages (*S. paradoxus SpCi* and *SpD*) were removed. Derived variants private to *S. paradoxus SpA, SpB* and *SpC*, present in at least 75% of strains were selected. Singletons and variants segregating at frequencies above 0.5 were removed to reduce the impact of misassignment of ancestral base. Mutation spectra were obtained from 34, 20321 and 3633 variants in *SpA, SpB* and *SpC*. A mutation spectrum for *S. cerevisiae* was obtained from a combined dataset of 1011 yeast genomes (Peter et al. 2018), with *S. paradoxus* genome (CBS432) used to assign ancestral alleles. Variants were selected with the same criteria for the combined set of all strains and the mutation spectrum was obtained from 944984 variants.

### Functional analysis of *de novo* mutations

The properties of *de novo* mutations were compared to null distributions expected if mutations arose in a random manner. The properties analyzed were whether mutations arise in coding or non-coding sequences, in essential or non-essential genes, and if they were synonymous or non-synonymous substitutions. For each cross and property, the null distribution accounted for the genomic composition of the parental lineages involved and the empirical mutation spectrum. For *S. paradoxus SpA, SpB* and *SpC* parents, the gene predictions, and annotations of representative strains (LL2012_001, MSH-604 and LL2011_012 respectively) from (Hénault et al. 2020) were used. For *S. cerevisiae*, genome annotations of the strain YPS128 from (Yue et al. 2017) were used. Coding sequences were defined as any sequence annotated as a protein-coding gene. Essential gene annotations were taken from the Saccharomyces Genome Deletion Project at http://www-sequence.stanford.edu/group/yeast_deletion_project/Essential_ORFs.txt. Synonymous and non-synonymous substitutions were scored by translating the sequence of annotated genes.

For each property, a null distribution was derived for each cross as follows. The GC content of coding vs. non-coding regions and essential vs. non-essential genes was calculated for the representative reference genome of each subgenome of a given cross. The GC contents were used to derive the probability that a substitution occurs in a non-coding or a non-essential region given that its initial state is an A/T or G/C base pair. Similarly, the codon frequency distribution of each reference subgenome was calculated for each cross. The per-codon count of possible synonymous and non-synonymous substitutions for each substitution type (A_C, A_G, A_T, C_A, C_G, C_T) was derived from the standard genetic code. The codon frequencies and substitution types counts per codon were used to derive the probability that a substitution is synonymous given its substitution type. Mutations were simulated to yield the same mutation count and spectrum as determined by the empirical *de novo* mutations for each cross. Mutations were sampled from a multinomial distribution corresponding to the mutation spectrum, and functional properties of each mutation were sampled from binomial distributions with the previously described probabilities. The random sampling procedure was repeated for 10000 iterations to produce distributions of functional properties (proportion of non-coding, non-essential and synonymous substitutions) from which 95% confidence intervals were derived.

### Analysis of DNA repair genes

In total, 202 genes present in either of the three assemblies used for mapping were selected including genes involved in DNA repair (GO:0006281), in DNA mismatch repair pathway (Phillips et al. 2021) and few other genes known to affect mutation rate or spectrum in yeast (Gou et al. 2019; Loeillet et al. 2020). Variant effects were annotated with SnpEff (Cingolani et al. 2012). We searched for derived variants relative to parental genotypes, present uniquely in one cross and segregating in high frequencies. To identify loss of genes through sequence deletions, we used previously identified copy number estimates in windows to identify significant copy number losses and gains (Wilcoxon Rank Sum test *P*-value <0.05) overlapping genes of interest. Amino Acid conservation scores and confidence intervals were obtained using the ConSurf server (Landau et al. 2005) together with a predicted protein structure.

## Data Availability

The data underlying this article are available in NCBI under project numbers: PRJNA515073 (short-sequence data) and PRJNA514804 (genome assemblies).

## Acknowledgments and funding information

We thank M. Drouin, J. Hallin, D. Biot-Pelletier and C. Bautista for their comments on the manuscript. C.R.L. holds the Canada Research Chair in Cellular Systems and Synthetic Biology. This work was supported by the Natural Sciences and Engineering Research Council of Canada (RGPIN-2020-04844 to C.R.L., Alexander Graham Bell doctoral scholarship to M.H. and G.C.), Genome Canada Grant (LSARP BIOSAFE 10106 to A.F.), Fonds de Recherche du Québec - Santé (post-doctoral scholarship to S.M.), and Fonds de recherche du Québec – Nature et technologies (doctoral scholarship to G.C.).

## Notes

### Competing Interest Statement

The authors have declared no competing interest.

## References

Bashir T, Sailer C, Gerber F, Loganathan N, Bhoopalan H, Eichenberger C, Grossniklaus U, Baskar R. 2014. Hybridization alters spontaneous mutation rates in a parent-of-origin-dependent fashion in Arabidopsis. Plant Physiol. 165:424–437.

Baym M, Kryazhimskiy S, Lieberman TD, Chung H, Desai MM, Kishony RK. 2015. Inexpensive multiplexed library preparation for megabase-sized genomes. PLoS One 10:1–15.

Boeva V, Popova T, Bleakley K, Chiche P, Cappo J, Schleiermacher G, Janoueix-Lerosey I, Delattre O, Barillot E. 2012. Control-FREEC: a tool for assessing copy number and allelic content using next-generation sequencing data. Bioinformatics 28:423–425.

Bolger AM, Lohse M, Usadel B. 2014. Trimmomatic: a flexible trimmer for Illumina sequence data. Bioinformatics 30:2114–2120.

Charron G, Marsit S, Hénault M, Martin H, Landry CR. 2019. Spontaneous whole-genome duplication restores fertility in interspecific hybrids. Nat. Commun. [Internet] 10. Available from: http://dx.doi.org/10.1038/s41467-019-12041-8

Cingolani P, Platts A, Wang LL, Coon M, Nguyen T, Wang L, Land SJ, Lu X, Ruden DM. 2012. A program for annotating and predicting the effects of single nucleotide polymorphisms, SnpEff: SNPs in the genome of Drosophila melanogaster strain w1118; iso-2; iso-3. Fly 6:80–92.

Deciphering Developmental Disorders Study. 2017. Prevalence and architecture of de novo mutations in developmental disorders. Nature 542:433–438.

Deem A, Keszthelyi A, Blackgrove T, Vayl A, Coffey B, Mathur R, Chabes A, Malkova A. 2011. Break-induced replication is highly inaccurate. PLoS Biol. 9:e1000594.

Demogines A, Wong A, Aquadro C, Alani E. 2008. Incompatibilities involving yeast mismatch repair genes: a role for genetic modifiers and implications for disease penetrance and variation in genomic mutation rates. PLoS Genet. 4:e1000103.

Drake JW. 1991. A constant rate of spontaneous mutation in DNA-based microbes. Proc. Natl. Acad. Sci. U. S. A. 88:7160–7164.

Eberlein C, Hénault M, Fijarczyk A, Charron G, Bouvier M, Kohn LM, Anderson JB, Landry CR. 2019. Hybridization is a recurrent evolutionary stimulus in wild yeast speciation. Nat. Commun. 10:923.

Ene IV, Farrer RA, Hirakawa MP, Agwamba K, Cuomo CA, Bennett RJ. 2018. Global analysis of mutations driving microevolution of a heterozygous diploid fungal pathogen. Proc. Natl. Acad. Sci. U. S. A. 115:E8688–E8697.

Garrison E, Marth G. 2012. Haplotype-based variant detection from short-read sequencing. arXiv [q-bio.GN] [Internet]. Available from: http://arxiv.org/abs/1207.3907

Gou L, Bloom JS, Kruglyak L. 2019. The Genetic Basis of Mutation Rate Variation in Yeast. Genetics 211:731–740.

Hénault M, Marsit S, Charron G, Landry CR. 2020. The effect of hybridization on transposable element accumulation in an undomesticated fungal species. Elife [Internet] 9. Available from: http://dx.doi.org/10.7554/eLife.60474

Hicks WM, Kim M, Haber JE. 2010. Increased mutagenesis and unique mutation signature associated with mitotic gene conversion. Science 329:82–85.

ICGC/TCGA Pan-Cancer Analysis of Whole Genomes Consortium. 2020. Pan-cancer analysis of whole genomes. Nature 578:82–93.

Jiang P, Ollodart AR, Sudhesh V, Herr AJ, Dunham MJ, Harris K. 2021. A modified fluctuation assay reveals a natural mutator phenotype that drives mutation spectrum variation within Saccharomyces cerevisiae. bioRxiv [Internet]:2021.01.11.425955. Available from: https://www.biorxiv.org/content/10.1101/2021.01.11.425955v1

Landau M, Mayrose I, Rosenberg Y, Glaser F, Martz E, Pupko T, Ben-Tal N. 2005. ConSurf 2005: the projection of evolutionary conservation scores of residues on protein structures. Nucleic Acids Res. 33:W299–W302.

Lang GI, Parsons L, Gammie AE. 2013. Mutation rates, spectra, and genome-wide distribution of spontaneous mutations in mismatch repair deficient yeast. G3 3:1453–1465.

Leducq J-B, Nielly-Thibault L, Charron G, Eberlein C, Verta J-P, Samani P, Sylvester K, Hittinger CT, Bell G, Landry CR. 2016. Speciation driven by hybridization and chromosomal plasticity in a wild yeast. Nat Microbiol 1:15003.

Li H. 2013. Aligning sequence reads, clone sequences and assembly contigs with BWA-MEM. arXiv [q-bio.GN] [Internet]. Available from: http://arxiv.org/abs/1303.3997

Li H. 2018. Minimap2: pairwise alignment for nucleotide sequences. Bioinformatics 34:3094–3100.

Liu H, Zhang J. 2019. Yeast Spontaneous Mutation Rate and Spectrum Vary with Environment. Curr. Biol. 29:1584–1591.e3.

Loeillet S, Herzog M, Puddu F, Legoix P, Baulande S, Jackson SP, Nicolas AG. 2020. Trajectory and uniqueness of mutational signatures in yeast mutators. Proc. Natl. Acad. Sci. U. S. A. 117:24947–24956.

Luria SE, Delbrück M. 1943. Mutations of Bacteria from Virus Sensitivity to Virus Resistance. Genetics 28:491–511.

Lynch M. 2007. The Origins of Genome Architecture. Sinauer Associates

Marsischky GT, Filosi N, Kane MF, Kolodner R. 1996. Redundancy of Saccharomyces cerevisiae MSH3 and MSH6 in MSH2-dependent mismatch repair. Genes Dev. 10:407–420.

Marsit S, Hénault M, Charron G, Fijarczyk A, Landry CR. 2021. The neutral rate of whole-genome duplication varies among yeast species and their hybrids. Nat. Commun. 12:1–11.

Ma X, Rogacheva MV, Nishant KT, Zanders S, Bustamante CD, Alani E. 2012. Mutation hot spots in yeast caused by long-range clustering of homopolymeric sequences. Cell Rep. 1:36–42.

Nguyen DT, Wu B, Long H, Zhang N, Patterson C, Simpson S, Morris K, Thomas WK, Lynch M, Hao W. 2020. Variable spontaneous mutation and loss of heterozygosity among heterozygous genomes in yeast. Mol. Biol. Evol. [Internet]. Available from: https://academic.oup.com/mbe/article-pdf/doi/10.1093/molbev/msaa150/33399075/msaa150.pdf

Peter J, De Chiara M, Friedrich A, Yue J-X, Pflieger D, Bergström A, Sigwalt A, Barre B, Freel K, Llored A, et al. 2018. Genome evolution across 1,011 Saccharomyces cerevisiae isolates. Nature 556:339–344.

Petrov DA, Schutzman JL, Hartl DL, Lozovskaya ER. 1995. Diverse transposable elements are mobilized in hybrid dysgenesis in Drosophila virilis. Proc. Natl. Acad. Sci. U. S. A. 92:8050–8054.

Phillips MA, Steenwyk JL, Shen X-X, Rokas A. 2021. Examination of gene loss in the DNA mismatch repair pathway and its mutational consequences in a fungal phylum. bioRxiv [Internet]:2021.04.13.439724. Available from: https://www.biorxiv.org/content/10.1101/2021.04.13.439724v1.full

Serero A, Jubin C, Loeillet S, Legoix-Né P, Nicolas AG. 2014. Mutational landscape of yeast mutator strains. Proc. Natl. Acad. Sci. U. S. A. 111:1897–1902.

Serrato-Capuchina A, Matute DR. 2018. The role of transposable elements in speciation. Genes [Internet] 9. Available from: http://dx.doi.org/10.3390/genes9050254

Sharp NP, Sandell L, James CG, Otto SP. 2018. The genome-wide rate and spectrum of spontaneous mutations differ between haploid and diploid yeast. Proc. Natl. Acad. Sci. U. S. A. 115:E5046–E5055.

Shor E, Fox CA, Broach JR. 2013. The yeast environmental stress response regulates mutagenesis induced by proteotoxic stress. PLoS Genet. 9:e1003680.

Stirling PC, Shen Y, Corbett R, Jones SJM, Hieter P. 2014. Genome destabilizing mutator alleles drive specific mutational trajectories in Saccharomyces cerevisiae. Genetics 196:403–412.

Swain PS, Stevenson K, Leary A, Montano-Gutierrez LF, Clark IBN, Vogel J, Pilizota T. 2016. Inferring time derivatives including cell growth rates using Gaussian processes. Nat. Commun. 7:13766.

Tattini L, Tellini N, Mozzachiodi S, D’Angiolo M, Loeillet S, Nicolas A, Liti G. 2019. Accurate Tracking of the Mutational Landscape of Diploid Hybrid Genomes. Mol. Biol. Evol. 36:2861–2877.

Venkataram S, Dunn B, Li Y, Agarwala A, Chang J, Ebel ER, Geiler-Samerotte K, Hérissant L, Blundell JR, Levy SF, et al. 2016. Development of a Comprehensive Genotype-to-Fitness Map of Adaptation-Driving Mutations in Yeast. Cell 166:1585–1596.e22.

Yang S, Wang L, Huang J, Zhang X, Yuan Y, Chen J-Q, Hurst LD, Tian D. 2015. Parent-progeny sequencing indicates higher mutation rates in heterozygotes. Nature 523:463–467.

Yue J-X, Li J, Aigrain L, Hallin J, Persson K, Oliver K, Bergström A, Coupland P, Warringer J, Lagomarsino MC, et al. 2017. Contrasting evolutionary genome dynamics between domesticated and wild yeasts. Nat. Genet. 49:913–924.

Zhu YO, Siegal ML, Hall DW, Petrov DA. 2014. Precise estimates of mutation rate and spectrum in yeast. Proc. Natl. Acad. Sci. U. S. A. 111:E2310–E2318.

