## Supplementary figures for "Heterogeneous mutation rates and spectra in yeast hybrids"

Supplementary Figure S1
Supplementary Figure S2
Supplementary Figure S3
Supplementary Figure S4
Supplementary Figure S5
Supplementary Figure S6
Supplementary Figure S7
Supplementary Figure S8
Supplementary Figure S9
Supplementary Figure S10
Supplementary Figure S11
Supplementary Figure S12
Supplementary Figure S13
Supplementary Figure S14
Supplementary Figure S15
Supplementary Figure S16
Supplementary Figure S17

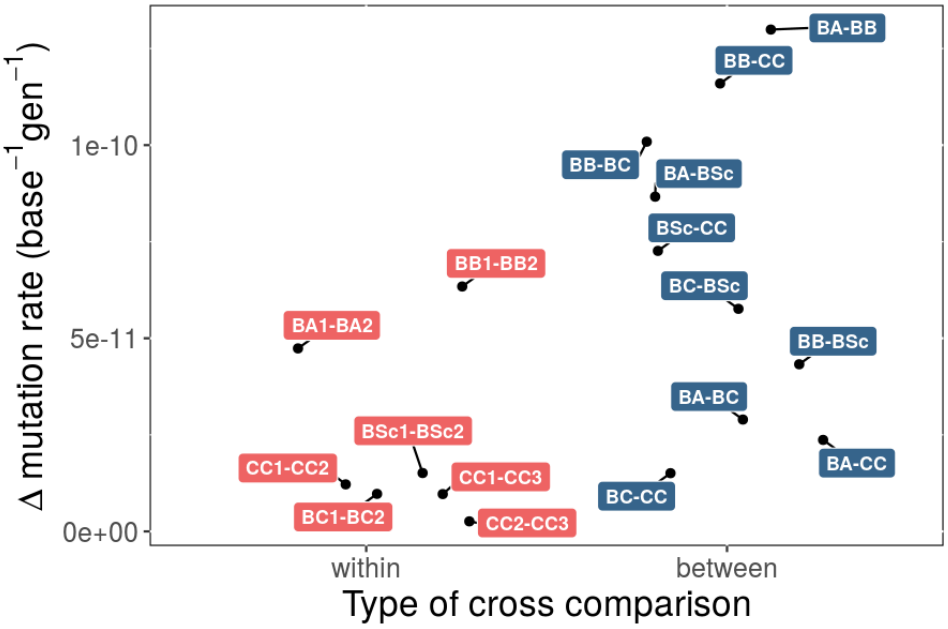


**Fig S1. Differences in mutation rates between pairs of crosses of the same type (within, red labels) overlap differences between pairs of crosses of different types (between, blue labels).** Highest differences belong to cross pairs including BB cross.

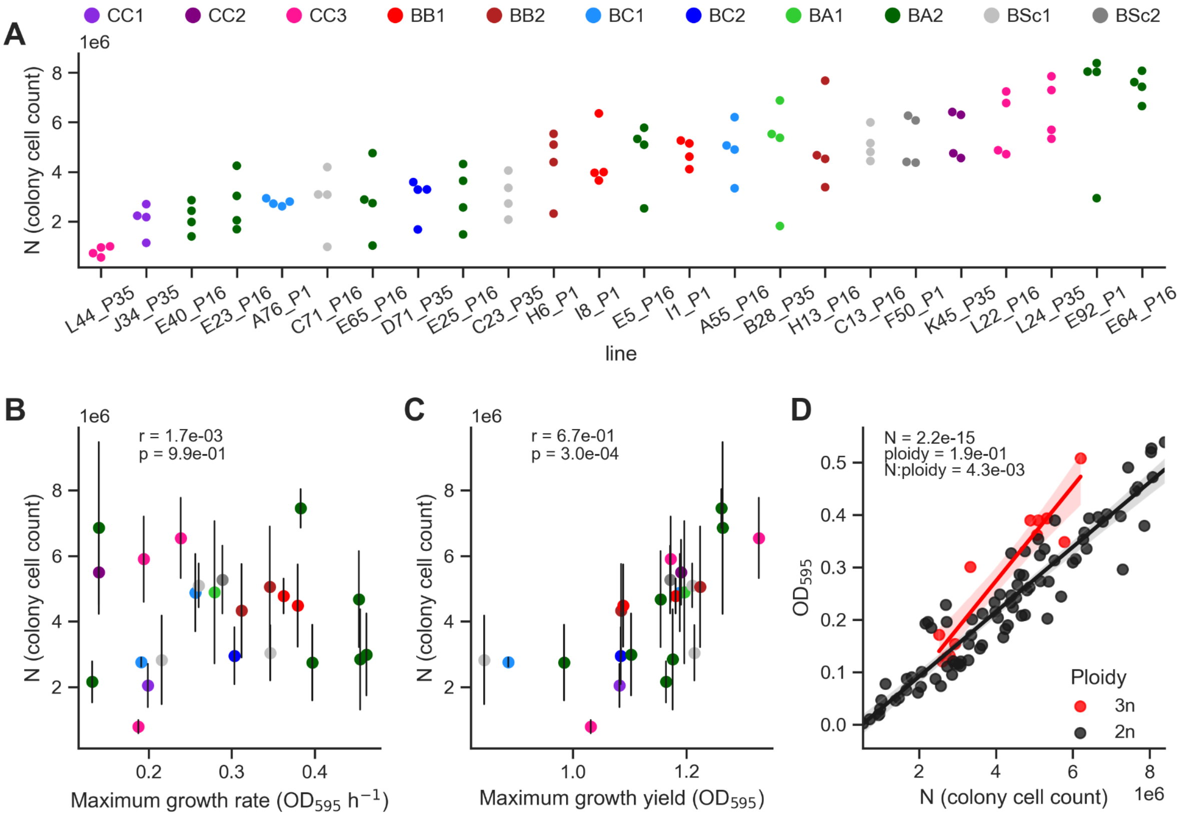


**Fig S2. Correlations between growth curve parameters and maximal population size during the MA experiment**. **A.** Cell counts per colony extrapolated from colony resuspension cell densities measured by flow cytometry. For 24 randomly selected MA lines, four colonies from independent streakings were analyzed. **B, C.** Pearson's correlation between colony cell count and maximum growth rate (**A**) or maximum growth yield (**B**) extracted from growth curves in liquid medium. Dots and lines show the mean and standard deviation in colony cell count, respectively. **D.** Ploidy has a significant effect on the optical density per cell. *P*-values are shown for the terms of a linear model fitted on $OD_{595}$ values.

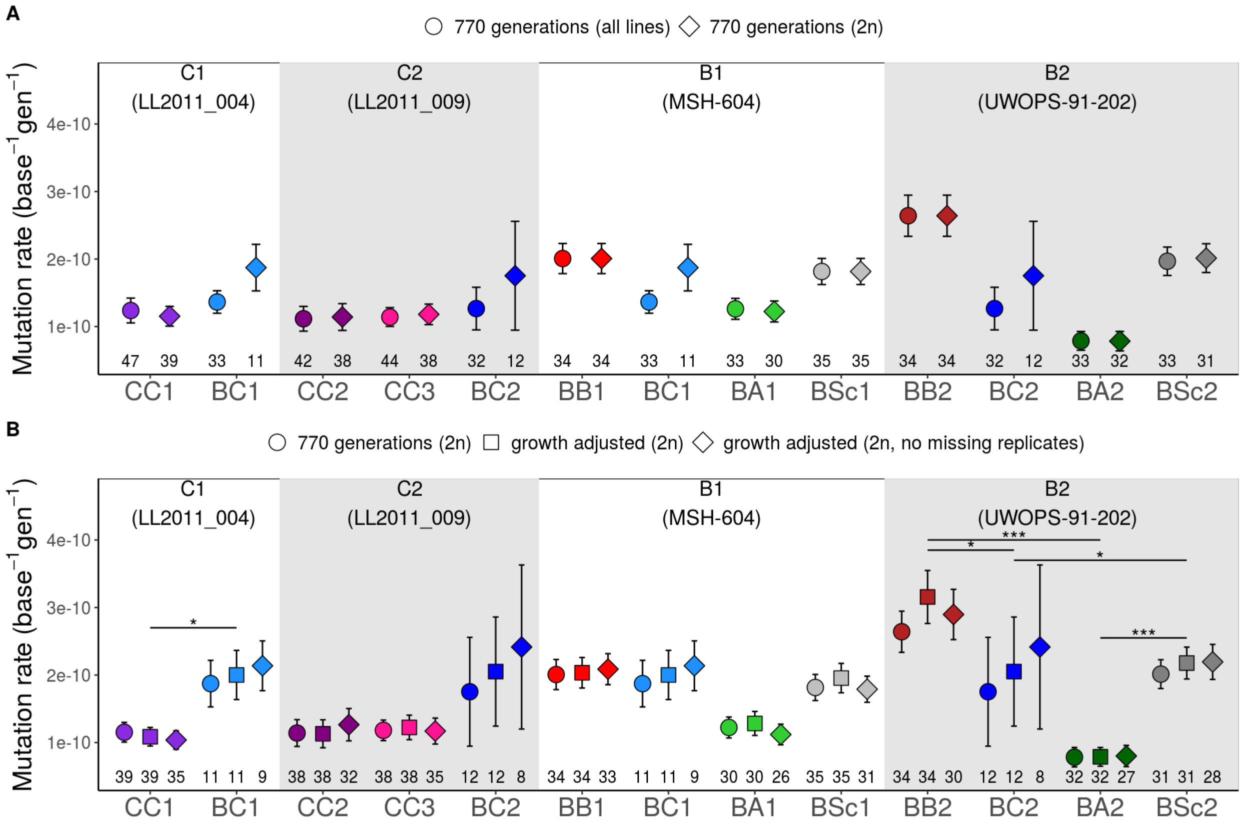


**Fig S3. Growth rate adjustment does not affect estimates of mutation rates.** Mean mutation rates per haploid position per generation with standard errors estimated from replicate lines shown in four groups sharing the same parental strain. **A.** Rates calculated with 770 generations for all lines (circles) and diploids only (diamonds). **B.** Rates calculated for diploid lines only with 770 generations (circles), number of generations adjusted for growth yield (square), and growth-adjusted rates with no missing replicates (diamonds). Mutation rates vary between crosses in C1 and B2 groups (rates calculated with 770 generations, Kruskal-Wallis test, *P*-values 0.033 and 2.04×10$^{-6}$ respectively). Asterisks show FDR-corrected differences at * *P*-value < 0.05, *** *P*-value < 0.0001, Wilcoxon Rank Sum Test. Number of lines is depicted at the bottom.

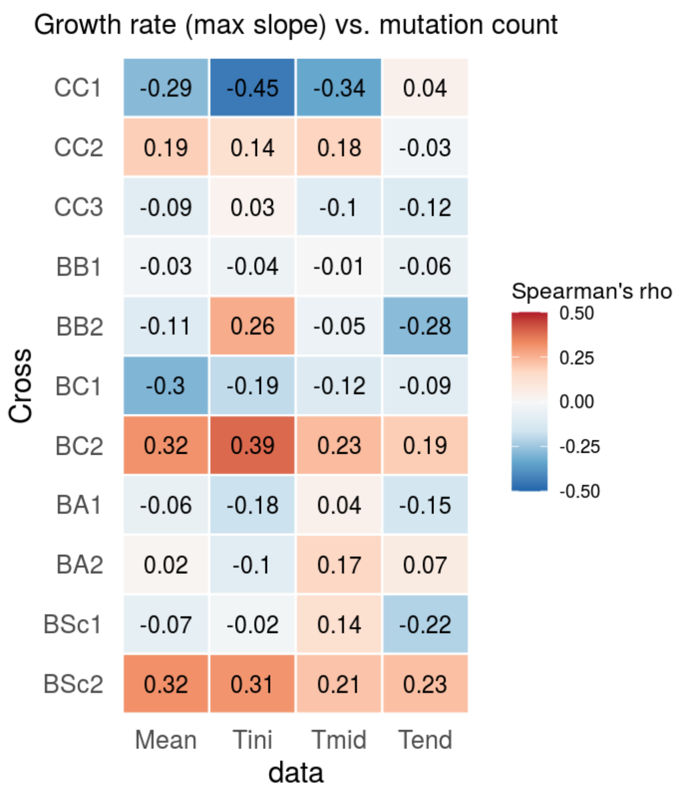


**Fig S4. Spearman's rank correlation coefficients between mutation counts ($5 \times 10^{-8}$) and growth rates for different crosses.** Growth yield calculated as a mean maximum yield (Mean) of 3 timepoints after one (Tini), 16 (Tmid) and 35 passages (Tend) for lines with no missing replicates. No correlations are significant after multiple test correction at FDR = 0.05.

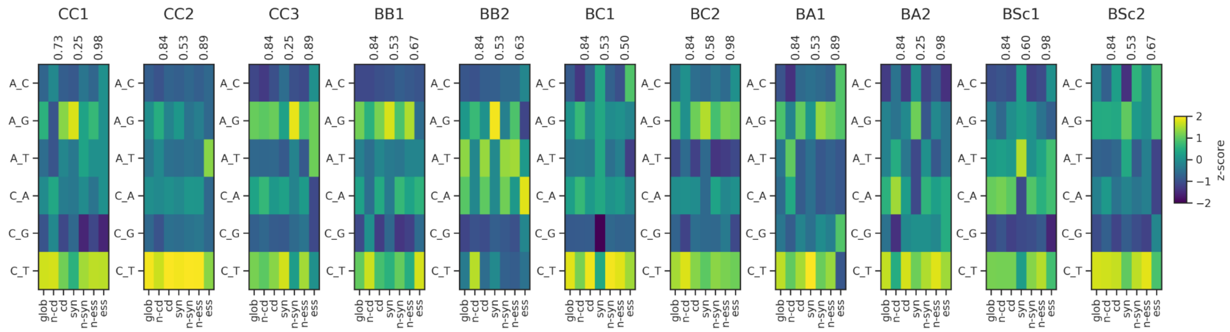


**Fig S5. Mutation spectra of individual crosses are similar across functional categories.** The z-score of mutation counts for each cross and functional category (leftmost being the global spectrum) is shown.

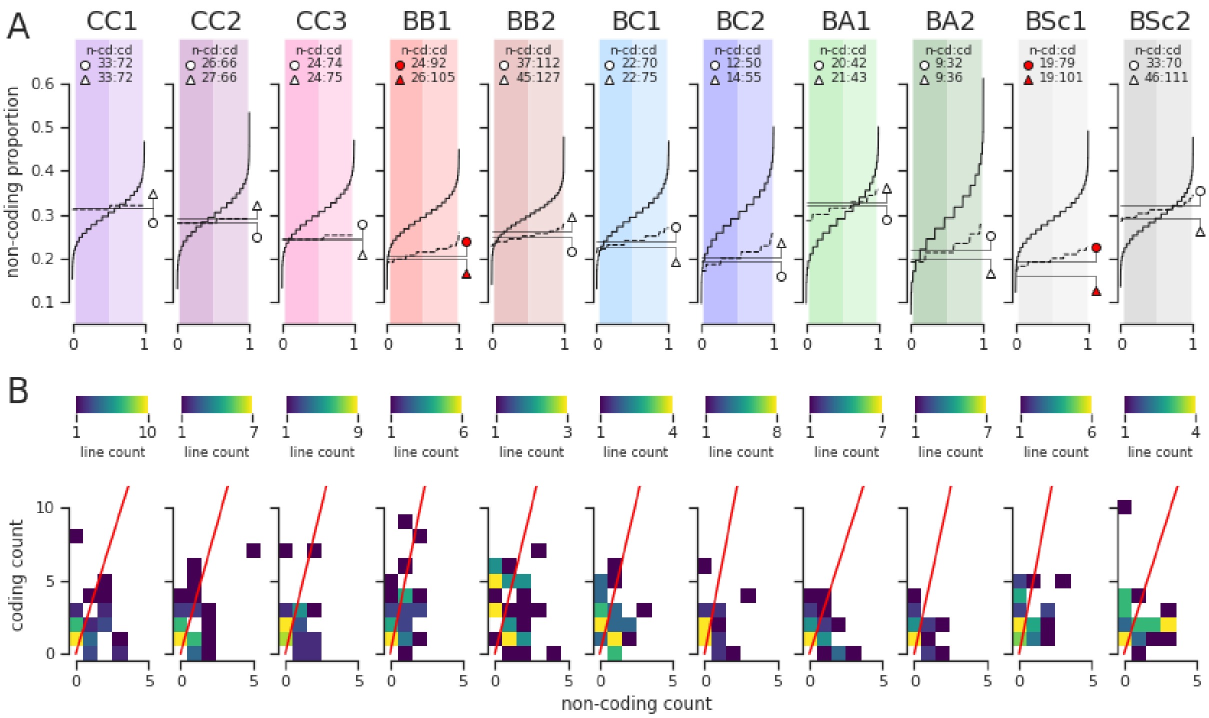

**Fig S6. Proportion of non-coding de novo mutations and neutral expectations for each cross. A.** The empirical proportion of non-coding mutations is shown by the solid horizontal lines with circular symbols. Solid horizontal lines with triangular symbols show the empirical proportion adjusted after including de novo mutations calls that were filtered out for overlapping with parental SNPs. Non-coding:coding ratios are detailed at the top of each graph. Solid line cumulative distributions show the null distribution of non-coding proportion for 10000 iterations of the random sampling procedure. Dashed line cumulative distributions show the empirical de novo mutations additioned with 10000 iterations of simulated mutations at loci overlapping with parental SNPs. Background shaded areas delimit the 0.025$^{th}$, 0.5$^{th}$ and 0.975$^{th}$ quantiles. Circular and triangular symbols are colored in red and blue when the point estimates fall below and above the 0.025$^{th}$ and 0.975$^{th}$ quantiles of the null distribution, respectively. **B.** Per-line counts of non-coding and coding de novo mutations. The red line shows the empirical ratio and the color map shows line counts.

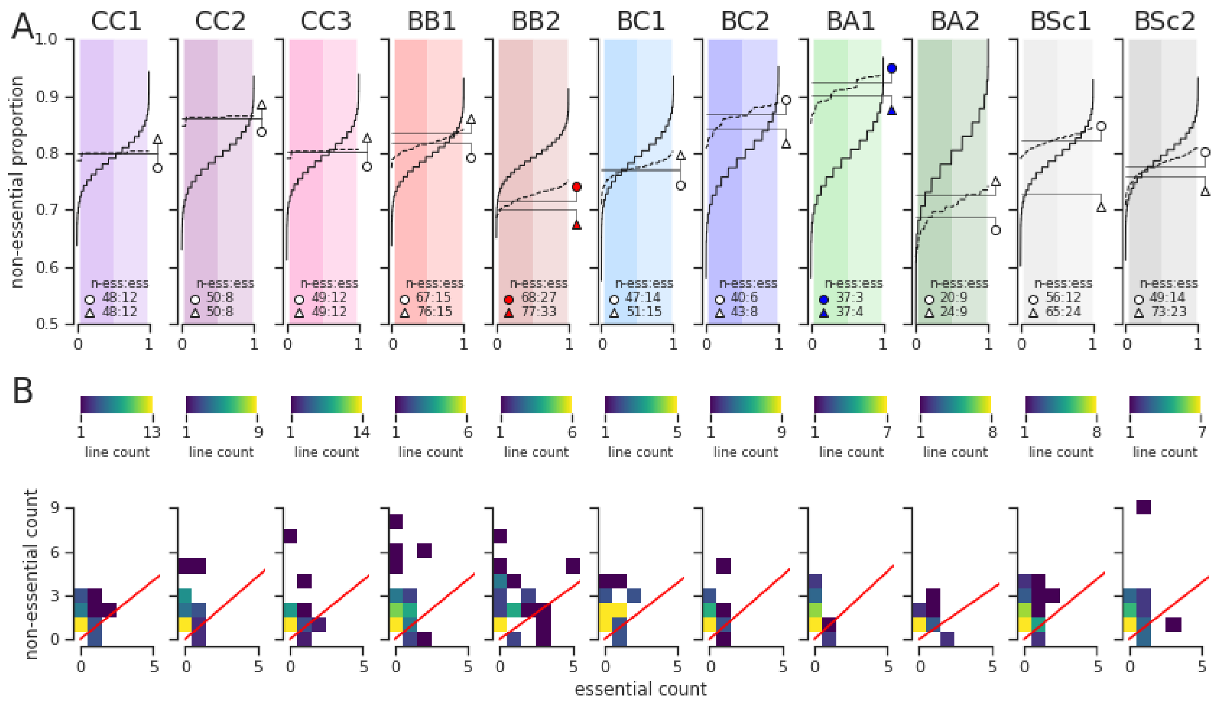


**Fig S7. Proportion of de novo mutations in non-essential genes and neutral expectations for each cross. A.** The empirical proportion of mutations in non-essential genes is shown by the solid horizontal lines with circular symbols. Solid horizontal lines with triangular symbols show the empirical proportion adjusted after including de novo mutations calls that were filtered out for overlapping with parental SNPs. Non-essential:essential ratios are detailed at the bottom of each graph. Solid line cumulative distributions show the null distribution of proportion of mutations in non-essential genes for 10000 iterations of the random sampling procedure. Dashed line cumulative distributions show the empirical de novo mutations additioned with 10000 iterations of simulated mutations at loci overlapping with parental SNPs. Background shaded areas delimit the $0.025^{th}$, $0.5^{th}$ and $0.975^{th}$ quantiles. Circular and triangular symbols are colored in red and blue when the point estimates fall below and above the $0.025^{th}$ and $0.975^{th}$ quantiles of the null distribution, respectively. **B.** Per-line counts of de novo mutations in non-essential and essential genes. The red line shows the empirical ratio and the color map shows line counts.

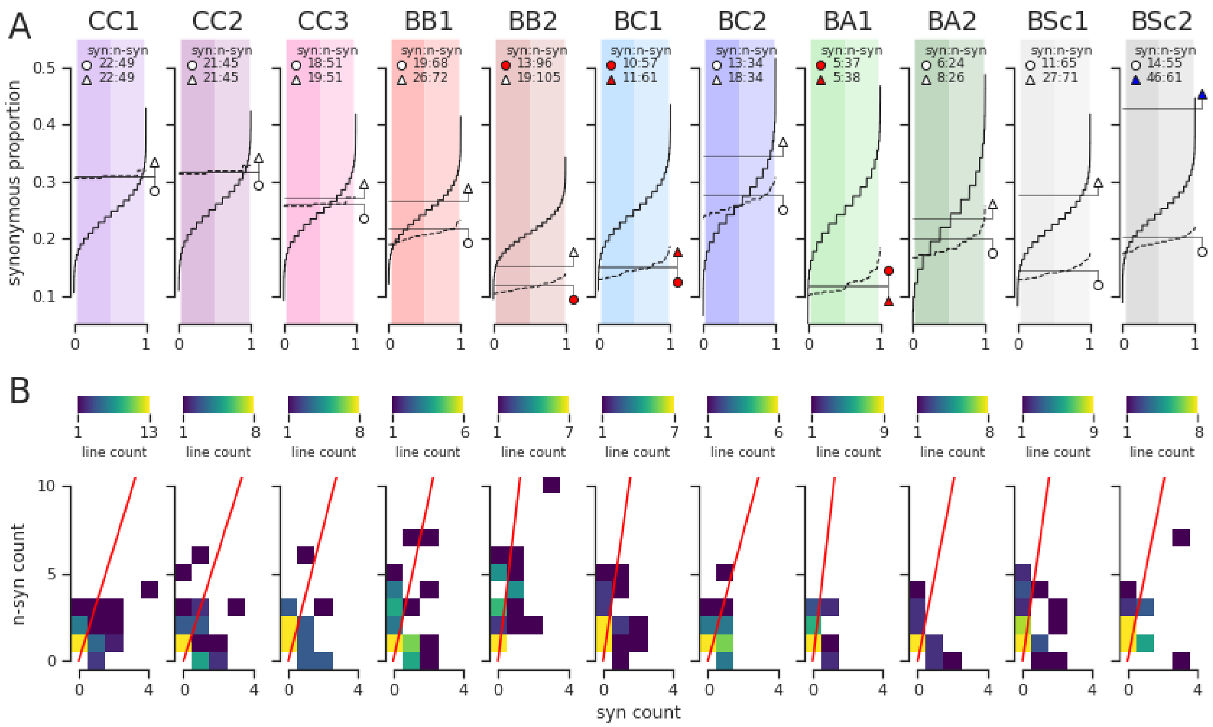


**Fig S8. Proportion of synonymous de novo mutations and neutral expectations for each cross. A.** The empirical proportion of synonymous mutations is shown by the solid horizontal lines with circular symbols. Solid horizontal lines with triangular symbols show the empirical proportion adjusted after including de novo mutations calls that were filtered out for overlapping with parental SNPs. Synonymous:non-synonymous ratios are detailed at the top of each graph. Solid line cumulative distributions show the null distribution of synonymous proportion for 10000 iterations of the random sampling procedure. Dashed line cumulative distributions show the empirical de novo mutations additioned with 10000 iterations of simulated mutations at loci overlapping with parental SNPs. Background shaded areas delimit the 0.025$^{th}$, 0.5$^{th}$ and 0.975$^{th}$ quantiles. Circular and triangular symbols are colored in red and blue when the point estimates fall below and above the 0.025$^{th}$ and 0.975$^{th}$ quantiles of the null distribution, respectively. **B.** Per-line counts of synonymous and non-synonymous de novo mutations. The red line shows the empirical ratio and the color map shows line counts.

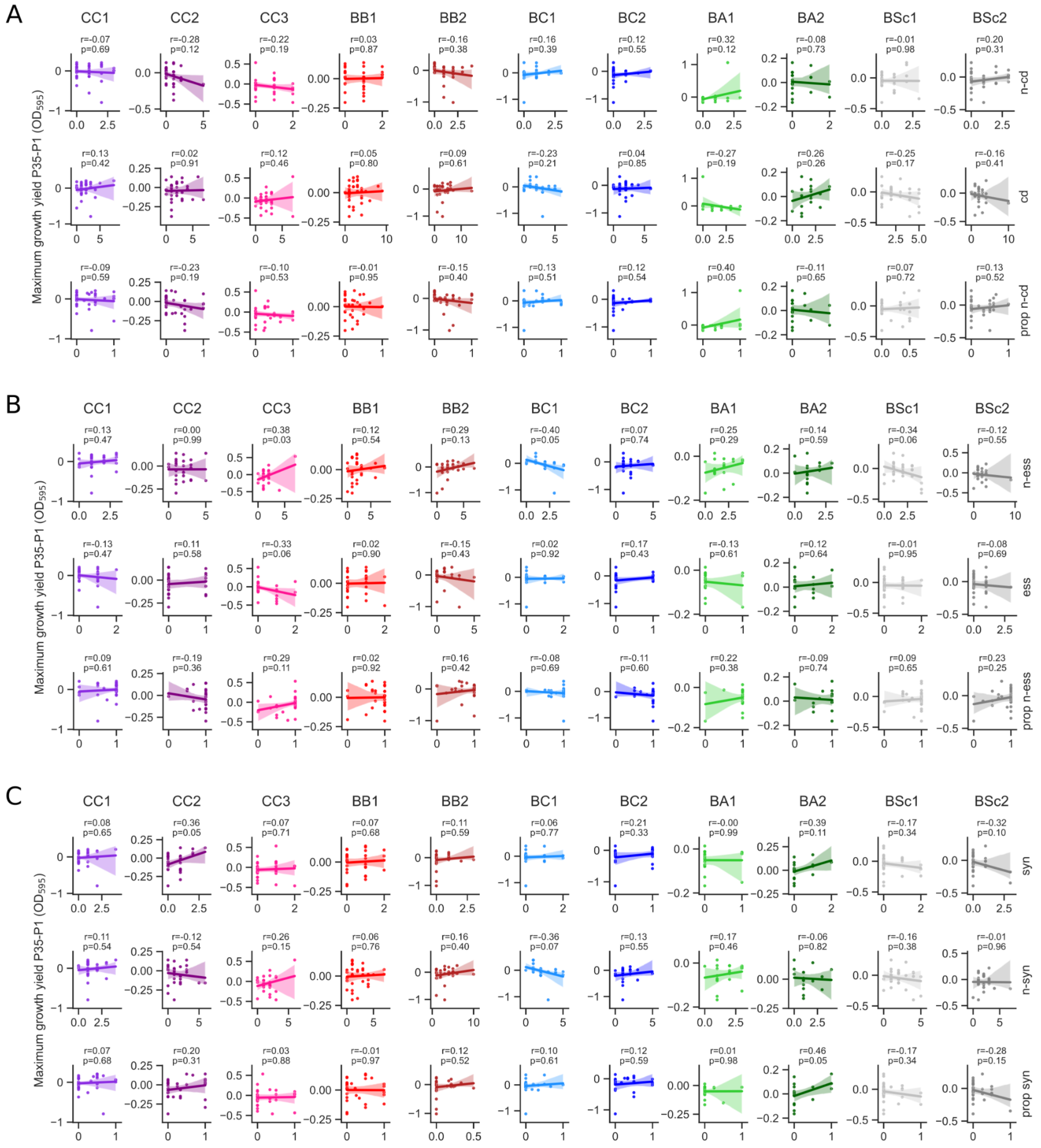


**Fig S9. Correlations of growth yield change with mutation counts, splitted by cross and functional category. A.** Counts of non-coding and coding mutations, and proportion of non-coding mutations. **B.** Counts of mutations in non-essential and essential genes, and proportion of mutations in non-essential genes. **C.** Counts of synonymous and non-synonymous mutations, and proportion of synonymous mutations. Pearson's r and *P*-value of individual linear regressions are shown on top of the plots.

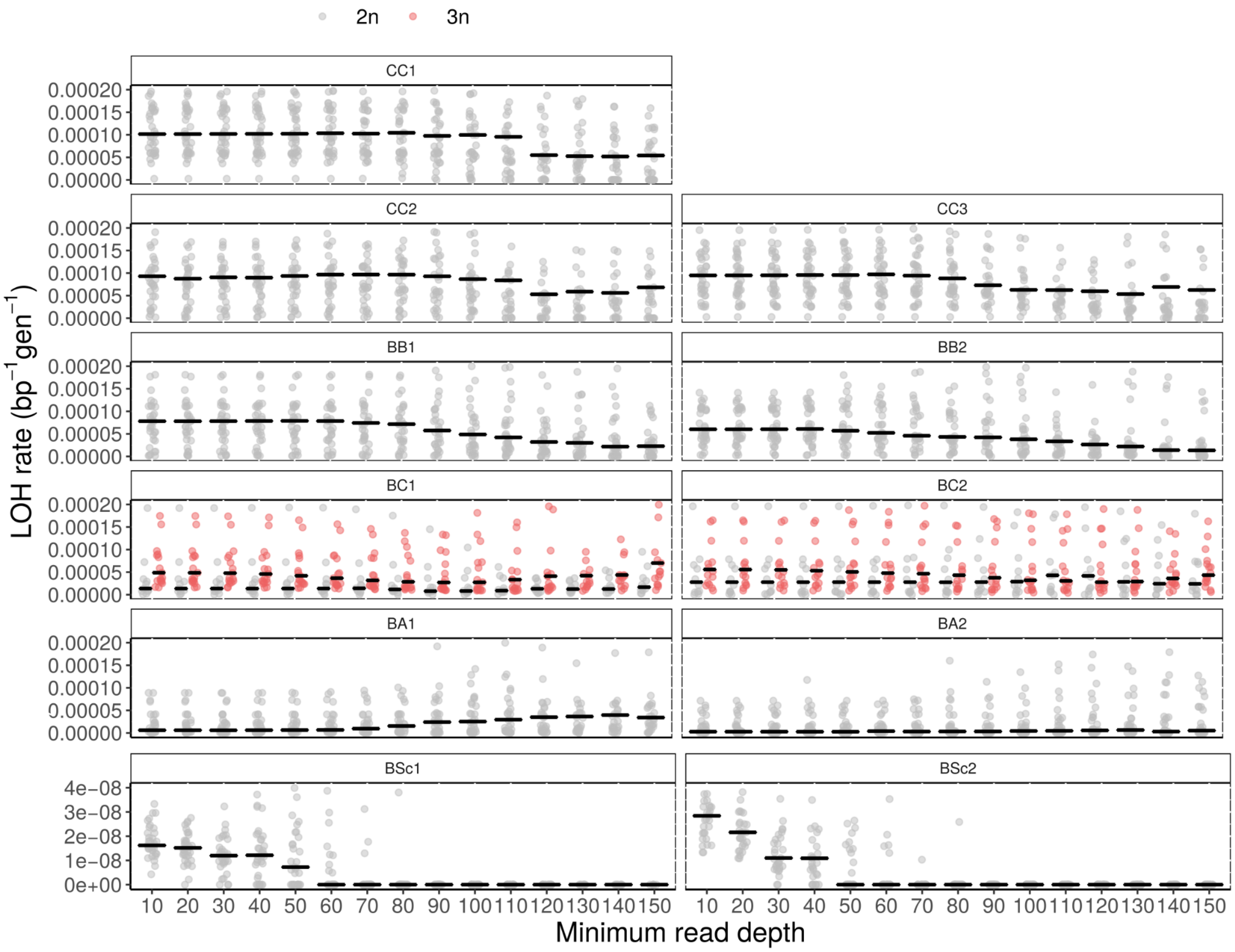


**Fig S10. Effect of minimum coverage on estimation of LOH in diploid and triploid lines.** Dots show estimated LOH rate calculated from the proportion of heterozygotes which had a changed genotype by $T_{end}$ in each line. Horizontal lines show medians. LOH rates increase for triploids in low coverage cutoff, therefore minimum read depth of 70 for CC1-BA2 crosses and minimum read depth of 40 for BSc crosses was considered for calculating LOH rates.

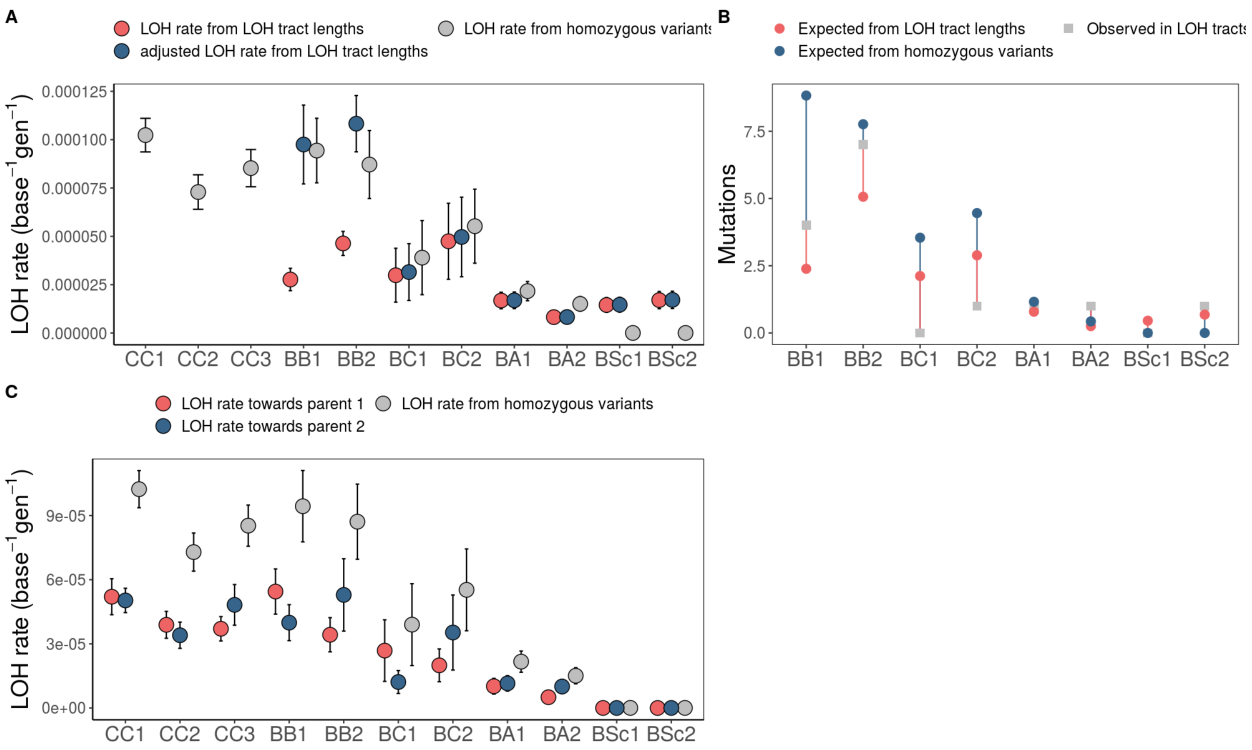


**Fig S11. Mean LOH rates estimated in diploid lines from LOH tract lengths** (Marsit et al. 2021) **and proportion of heterozygotes turned into homozygotes by $T_{end}$. A.** Comparison of LOH rates calculated from LOH tract lengths (red circles), rates calculated from LOH tract lengths but corrected by the probability of detecting a 1000 bp window with 10 heterozygous variants (minimum for calling an LOH tract, blue circles), and rates calculated from the proportion of homozygous variants (grey circles). **B.** Comparison of the total number of *de novo* mutations found within LOH tracts (grey squares), with the number of expected mutations given LOH lengths (red circles), and the number of expected mutations given the proportion of homozygous variants (blue circles). **C.** LOH rate towards the first parent in the cross (red circle) or the second parent in the cross (blue circle).

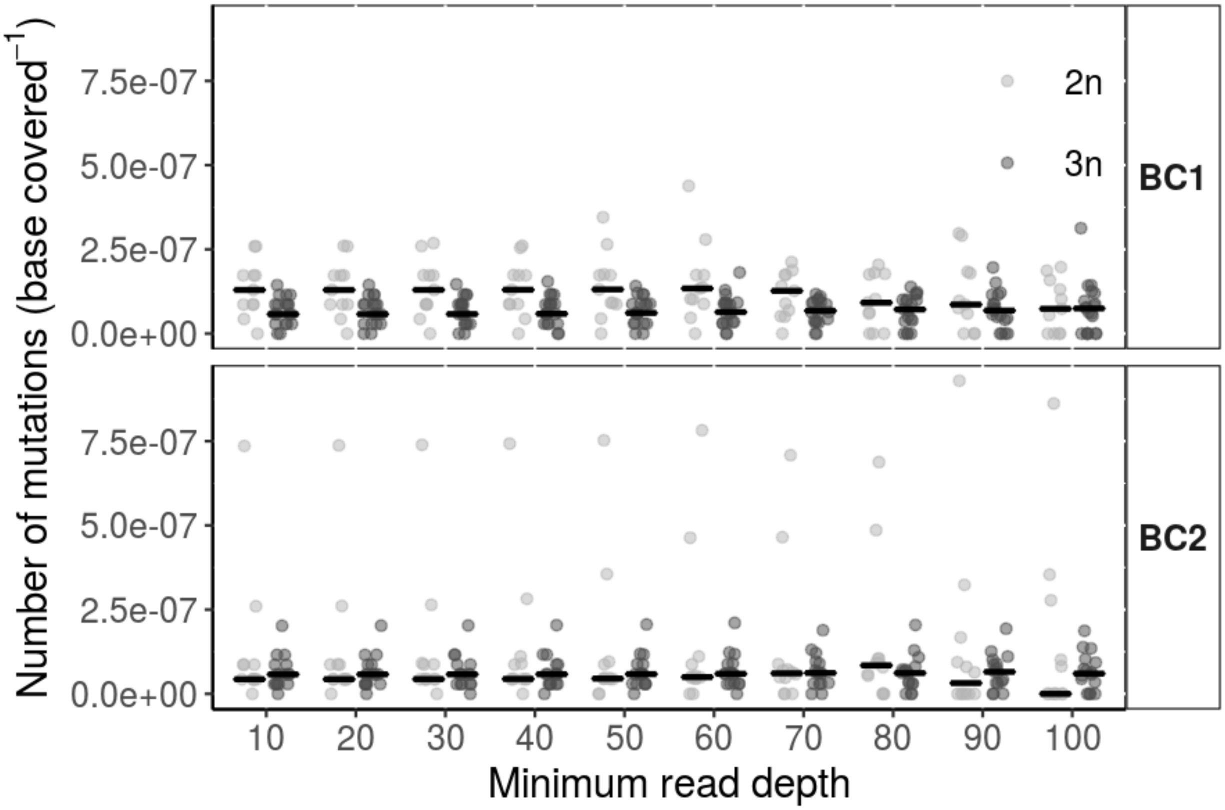


**Fig S12. Mutation rate differences between diploid and triploid lines from BC crosses.** Mutation rates in diploid and triploid lines after removing low covered bases from the genome. Lines indicate medians, dots correspond to raw mutation counts divided by the number of positions in the genome with a given depth and polyploidy.

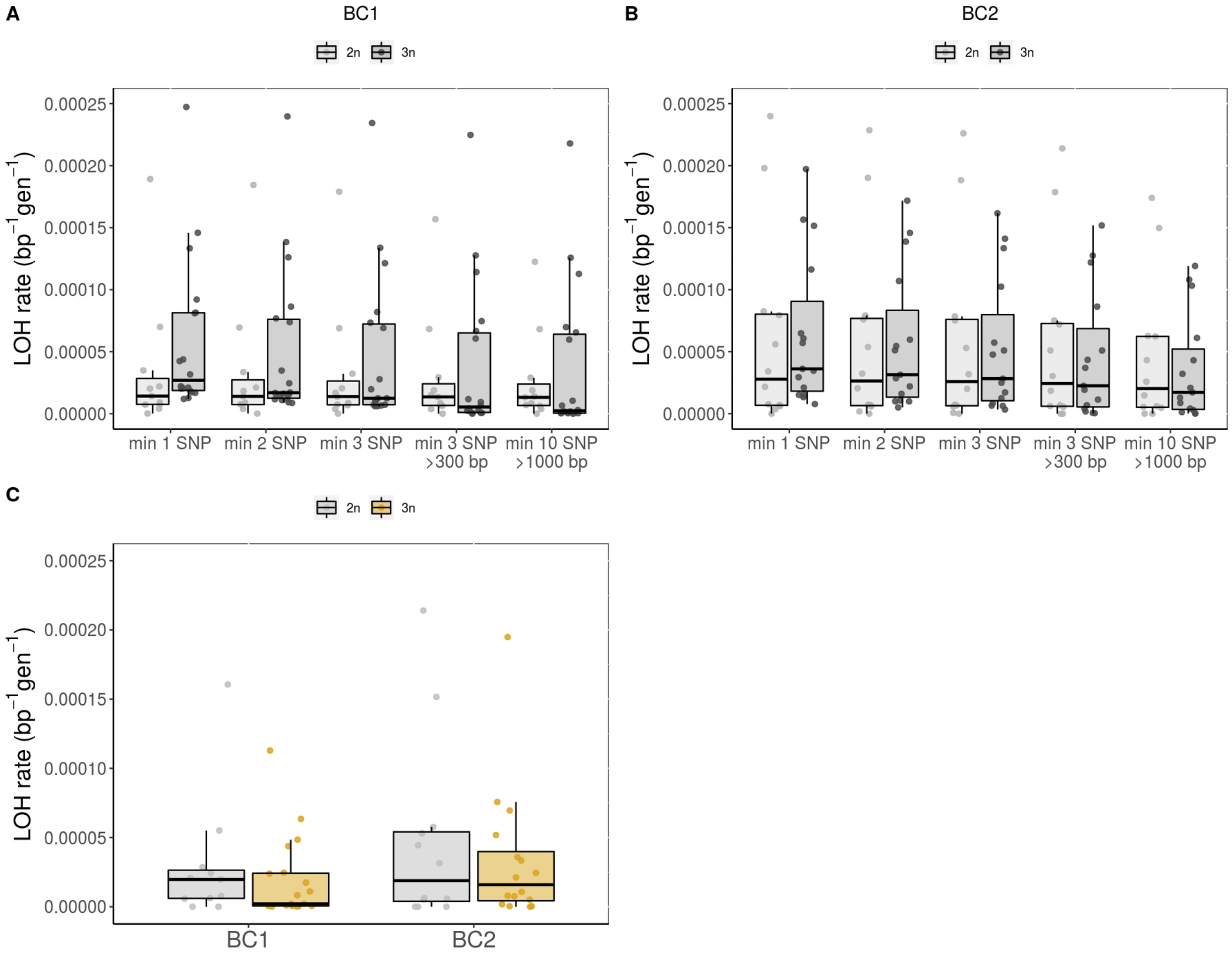


**Fig S13. Impact of different filtering criteria on LOH rate calculation in diploid and triploid BC crosses**. LOH rates with different filtering criteria in diploids and triploids in BC1 (**A**) and BC2 (**B**) cross. LOH rates were calculated from heterozygous positions with changed genotypes without any filtering (min 1 SNP, same as fig. 2B), including only tracts of at least 2 consecutive SNPs (min 2 SNP), tracts of at least 3 SNPs (min 3 SNPs), tracts of 3 at least SNPs and min 300 bp long (min 3 SNP > 300 bp), and tracts of at least 10 SNPs and min 1000 bp long (min 10 SNP > 1000 bp). **C.** LOH rate calculated from LOH tracts of minimum 1000 bp (and min 3 SNPs in 300bp) from (Marsit et al. 2021).

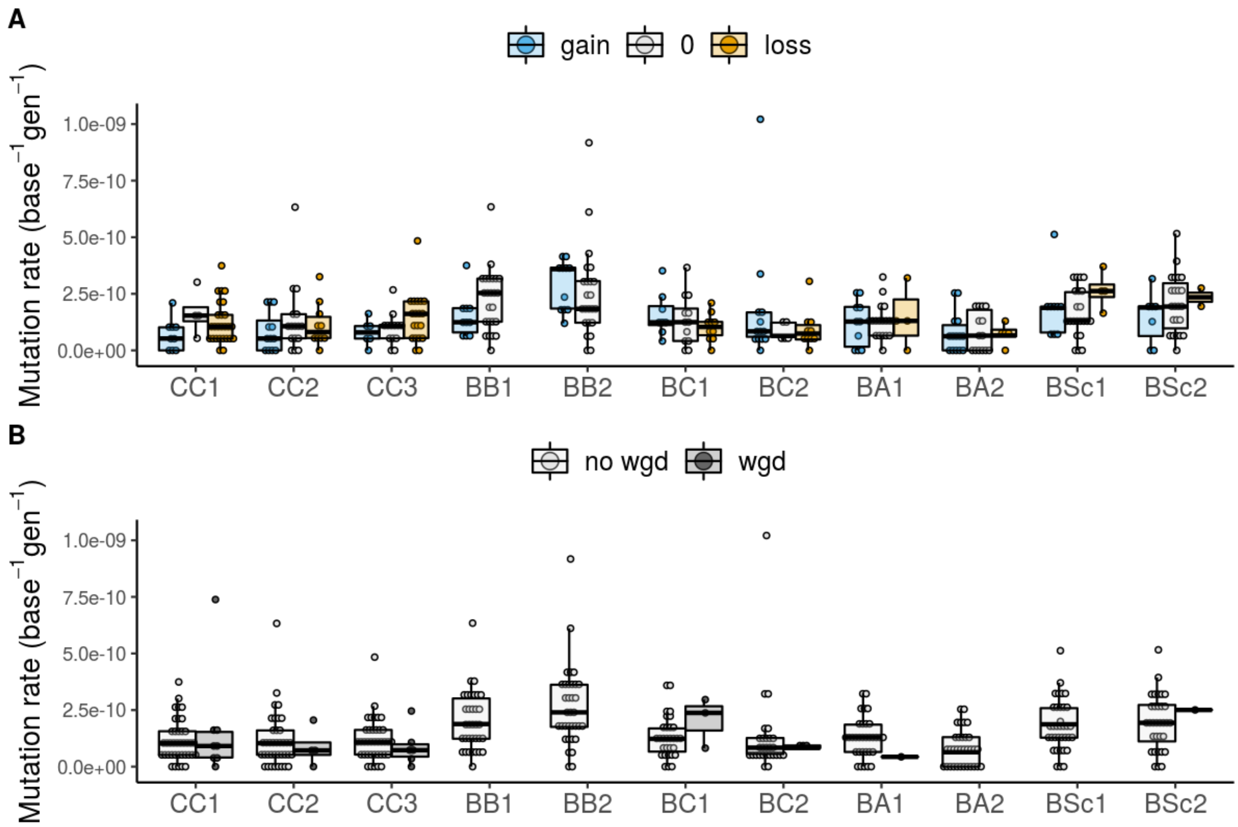


**Fig S14. Comparison of mutation rates between euploid lines and lines with aneuploidies or whole genome duplications. A**. Comparison of mutation rates in diploid and triploid lines, which lost (blue) or gained (orange) chromosomes in the course of experiment. 0 stands for no change of chromosomes. **B**. Comparison of mutation rates between diploid lines (no wgd) and lines which underwent whole genome duplication (wgd) in the course of experiment. Wilcoxon rank sum test of lower mutation rate for lines with gains of chromosomes or whole genome duplication not significant after multiple test correction at FDR = 0.05.

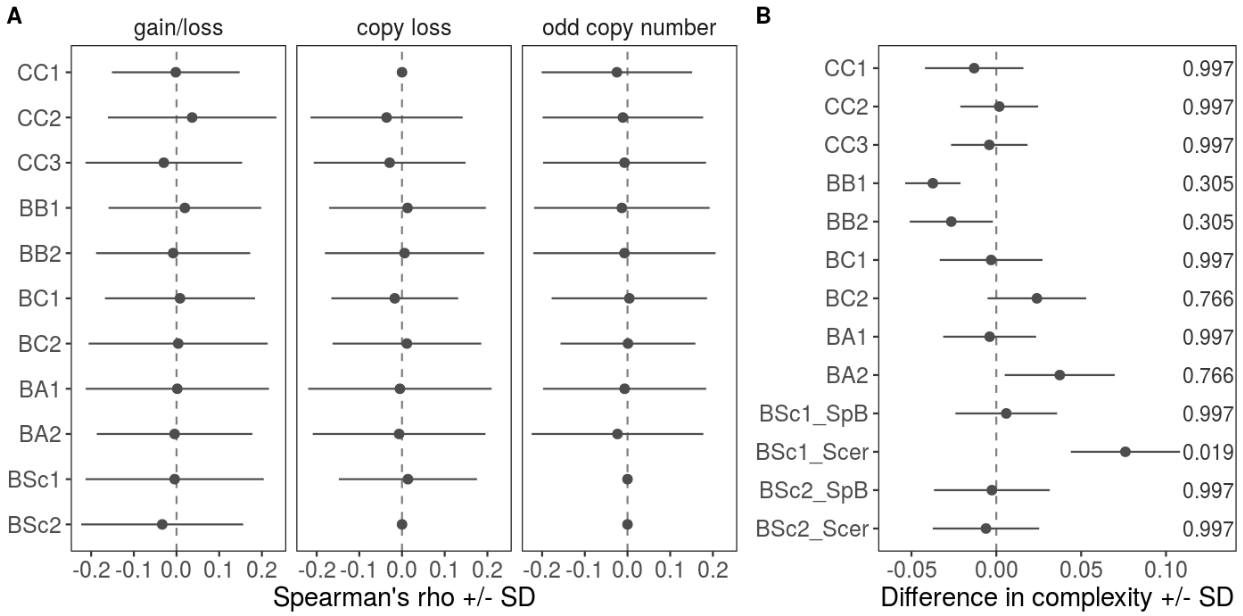


**Fig S15. Characteristics of DNA repair gene copies and sequence complexity among crosses. A.** No significant relationship between number of mutations and dynamics of copies of DNA repair genes. Points correspond to mean Spearman's rho between mutation count and a given measure per line with standard deviation after 100 bootstraps. Measures include number of significant gains/losses of copy number compared to background ploidy (first panel), number of genes with copy loss compared to expected diploid number regardless of background ploidy (second panel), and the number of genes with odd number of copies (third panel). **B.** Mean differences of median sequence complexity around *de novo* and random mutations with standard deviation after 100 bootstraps. Complexity was calculated for a sequence +/- 10bp around the mutation. Random mutations (in the same number as *de novo*) were sampled from the reference genomes excluding repeats and positions differentiating the parents, and according to the cross mutation spectrum. BH-adjusted *P*-values (at FDR = 0.05) from Mann-Whitney U test are shown to the right. In BSc crosses mutations were sampled separately on *S. paradoxus SpB* (_SpB) and *S. cerevisiae* (_Scer) genomes.

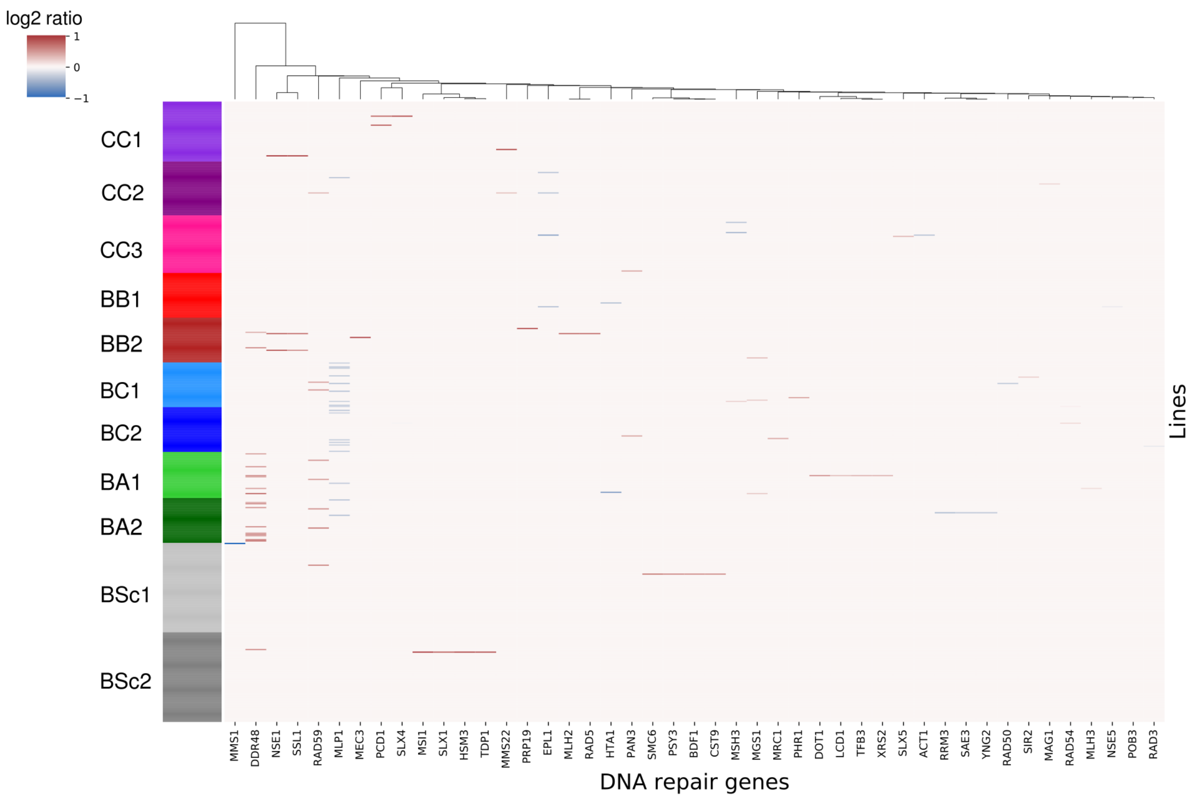


**Fig S16. No gene losses in DNA repair genes in BB2 at the initial timepoint of MA.** Heatmap shows significant gene losses and gains compared to chromosome background ploidy at the initial timepoint of MA experiment. Log2 ratio of 1 indicates doubling and -1 indicates loss of half of the copies compared to background copy number.

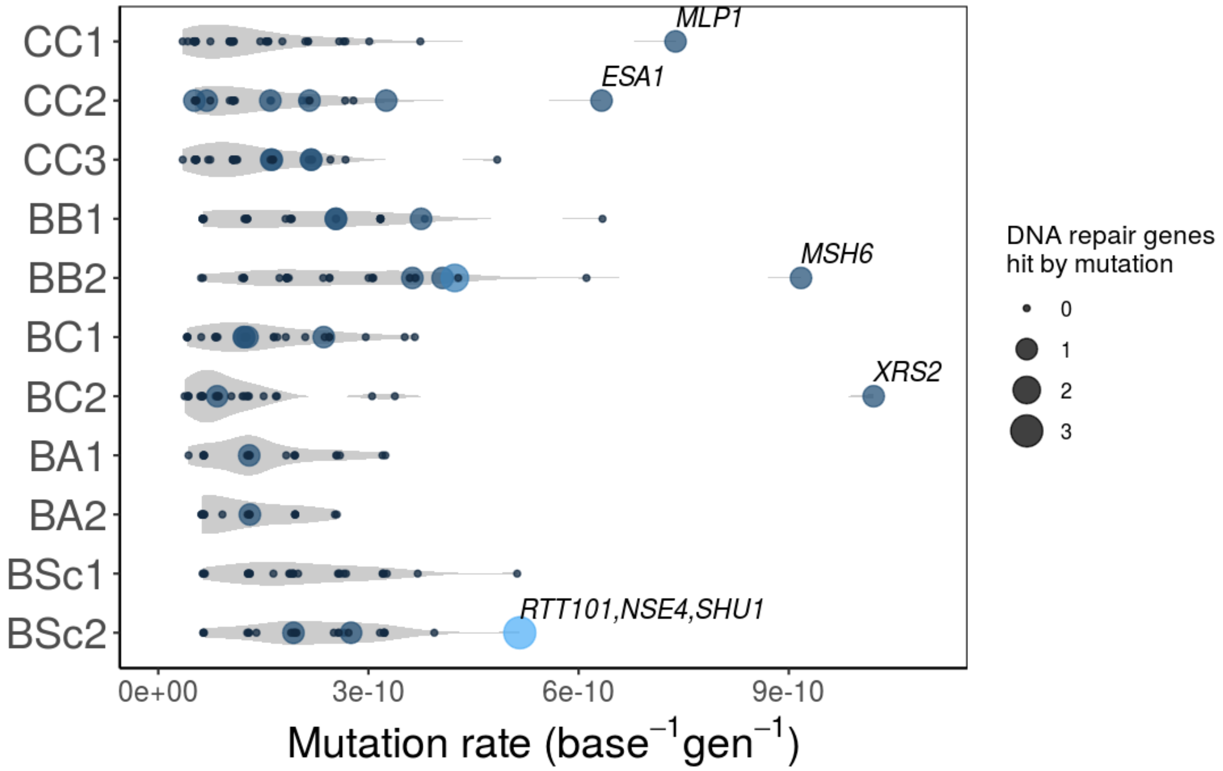


**Fig S17. Presence of non-synonymous or non-sense *de novo* mutations in DNA repair genes in MA lines.** Mutation rates of individual lines are shown, with dot size indicating the number of DNA repair genes hit by a non-synonymous/non-sense mutation. Lines with outlier mutation rates are labelled with the corresponding DNA repair genes.

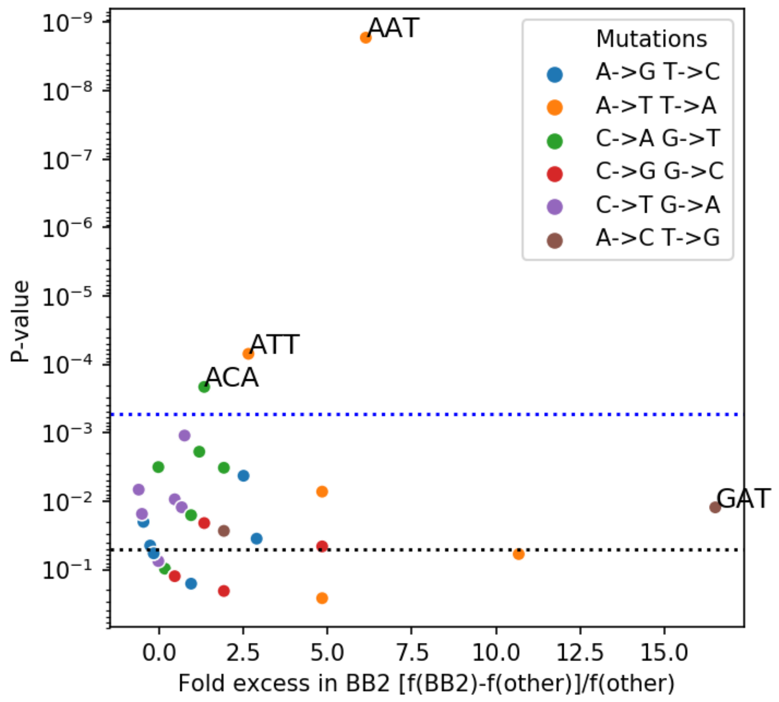


**Fig S18. Frequency of mutation changes in BB2 compared to other crosses in the context of surrounding nucleotides.** Each dot represents a mutated position with two surrounding nucleotides. Differences were tested using the chi-square test. The blue dotted line indicates *P*-value after Bonferroni correction, and black dotted line *P*-value = 0.05.
